# Characterization of *Arabidopsis thaliana* promoter bidirectionality and antisense RNAs by depletion of nuclear RNA decay enzymes

**DOI:** 10.1101/809194

**Authors:** Axel Thieffry, Jette Bornholdt, Maxim Ivanov, Peter Brodersen, Albin Sandelin

**Affiliations:** Department of Biology, University of Copenhagen, Ole Maaløes Vej 5, DK2200 Copenhagen N, Denmark; Biotech Research and Innovation Centre, University of Copenhagen, Ole Maaløes Vej 5, DK-2200 Copenhagen N, Denmark; Department of Plant and Environmental Sciences, University of Copenhagen, Thorvaldsensvej 40, DK-1871 Frederiksberg C, Denmark

## Abstract

In animals, transcription by RNA polymerase II initiates bidirectionally from gene promoters to produce pre-mRNAs on the forward strand and promoter upstream transcripts (PROMPTs) on the reverse strand. PROMPTs are rapidly degraded by the nuclear exosome. Similarly, active enhancer regions in animals initiate transcription of exosome-sensitive enhancer RNAs (eRNAs). Previous studies based on nascent RNA approaches concluded that *Arabidopsis thaliana* does not produce PROMPTs. Here, we used steady-state RNA sequencing methods in mutants defective in nuclear RNA decay, including by the exosome, to reassess the existence of PROMPTs and eRNAs in *A. thaliana*. While PROMPTs are overall rare in *A. thaliana*, about 100 clear cases of exosome-sensitive PROMPTs and 113 loci producing eRNA-like transcripts were identified. In addition, we found ∼200 transcription start sites within 3’-UTR-encoding regions that produce unspliced exosome-sensitive antisense RNAs covering much of the cognate pre-mRNA. A typical representative of this class of RNAs is the previously characterized non-coding RNA controlling the expression of the key seed dormancy regulator, *DELAY OF GERMINATION1*. Exosome-sensitive antisense RNAs are overrepresented in transcription factor genes, suggesting a potential for widespread control of gene expression. Lastly, we assess the use of alternative promoters in *A. thaliana* and compare the accuracy of existing TSS annotations.

## INTRODUCTION

The vast majority of promoters of mammalian protein-coding genes initiates transcription by RNA polymerase II on both strands, producing a sense pre-mRNA transcript and a shorter, unstable RNA on the reverse strand, denominated promoter upstream transcript (PROMPT) or upstream antisense RNA (uaRNA) (Core et al., 2008; Seila et al., 2008; Preker et al., 2008). Initiation of PROMPT and pre-mRNA transcription occurs at the edge of the nucleosome depleted region (NDR) in a bidirectional manner (Ntini et al., 2013; Andersson et al., 2014a, 2014b; Dubbury et al., 2018). Unlike pre-mRNAs, PROMPTs are rapidly degraded by the nuclear exosome (Preker et al., 2008), a central player in RNA degradation. The core exosome is a highly conserved 9-subunit complex consisting of a central, hexameric barrel (a trimer of RRP41-RRP42, RRP45-RRP46 and RRP43-MTR3 heterodimers), and a 3-subunit lid containing the proteins RRP4, RRP40, and CSL4. 3’-5’ exonucleases associate with the core exosome, and the resulting exosome machinery constitutes the main 3’-5’ exonuclease activity both in the nucleus and in the cytoplasm (Chlebowski et al., 2013; Kilchert et al., 2016). The exosome sensitivity of PROMPTs correlates with the frequency of occurrence of specific DNA sequence elements downstream of the PROMPT transcription start sites (TSS): compared to the coding strand, the PROMPT region is enriched in TSS-proximal polyadenylation sites and depleted of splice donor sites (Almada et al., 2013; Core et al., 2014; Ntini et al., 2013).

The pattern of bidirectional transcription initiation is not exclusive to promoters of coding genes, but appears to be generic for RNA polymerase II initiation in metazoans: it is also detectable at long noncoding RNA (lncRNA) promoters (Andersson et al., 2014b) and in particular at active enhancers (Kim et al., 2010). We and others have previously shown that enhancer regions can be accurately predicted as sites of bidirectional initiation of unstable transcripts on the basis of either RNA-seq (Wu et al., 2014), sequencing of capped RNA 5’-ends (Andersson et al., 2014a), or nascent RNA approaches (Core et al., 2014).

The similarity of transcription initiation patterns between different RNA classes has spurred the hypothesis that RNA polymerase II is subject to the same rules regardless of the RNA that is produced, so that bidirectionally transcribed NDRs constitute a generic transcription initiation block, and the fate and function of RNAs produced are determined after transcription initiation (Chen et al., 2017; Andersson et al., 2015b), also reviewed in (Andersson and Sandelin, 2019). It is currently an open question whether this framework is generic across eukaryotes. Recently, studies showed that the majority of enhancers and a substantial fraction of gene promoters are divergently transcribed in *Drosophila melanogaster* cells (Rennie et al., 2018) and larvae (Meers et al., 2018). In yeast species, the number of enhancers is small due to their compact genome. Promoter bidirectionality was observed in *Saccharomyces cerevisiae* (reviewed in (Jensen et al., 2013)), and in at least a subset of promoters in *Schizosaccharomyces pombe* (Shetty et al., 2017; Thodberg et al., 2018).

In plants, only a handful of studies has investigated this question in depth, using approaches in which nascent RNA is either labeled and affinity-purified from isolated nuclei (Global Run-On sequencing, GRO-seq (Core et al., 2008) and GRO-cap), or isolated as RNA polymerase II-associated RNA by immunoprecipitation (Native Elongating Transcript sequencing, NET-seq). Analysis of *Arabidopsis thaliana* seedlings by GRO-seq provided little support for bidirectional promoter transcription (Hetzel et al., 2016), and metaplots of GRO-seq and NET-seq signals also led Zhu et al. to conclude that promoters of protein-coding genes in *Arabidopsis* are unidirectional (Zhu et al., 2018). One study on maize also found little evidence for bidirectional transcription using GRO-seq (Erhard et al., 2015), but a precision nuclear run-on sequencing (PRO-seq) analysis of cassava and a re-analysis of the maize GRO-seq data generated by Erhard et al. showed many instances of clear bidirectional transcription at enhancer candidates (Lozano et al., 2018). In addition, this study identified some cases of bidirectionally transcribed protein-coding gene promoters in cassava (Lozano et al., 2018). Although both GRO-seq and NET-seq data clearly identify RNA populations different from the steady-state pool, a number of concerns may be raised on negative findings based on these techniques. First, GRO-seq can have limited power to detect weakly transcribed loci such as PROMPTs if the sequencing depth is low (Andersson et al., 2015a), and indeed, many of the published plant GRO-seq studies suffer from a lack of sample replicates. Second, if the degradation of particular RNA species is rapid and occurs co-transcriptionally, such RNAs may escape detection, even if the techniques successfully capture many other species of nascent RNA. For these reasons, it is not completely clear whether promoters in plants, including *A. thaliana*, truly are unidirectional, in particular because conflicting conclusions have been reached in maize using these techniques (Erhard et al., 2015; Lozano et al., 2018). The extent of bidirectional transcription in other genomic regions than gene promoters is similarly a question worthy of further investigation.

Although nascent RNA techniques also support bidirectional transcription at gene promoters in mammals (Andersson et al., 2015a), one of the most instrumental techniques in uncovering this aspect of RNA polymerase II function have been analysis of steady-state RNA in reference cells compared to cells in which components of the nuclear RNA exosome machinery were knocked down (Preker et al., 2008; Ntini et al., 2013). In *A. thaliana*, stable knockout mutants of the core exosome subunits RRP4 and RRP41 are not viable, while, surprisingly, the knockout of the third lid subunit CSL4 displays mild, if any, growth phenotype, and only a few molecular phenotypes (Chekanova et al., 2007). These findings originally inspired Chekanova et al. to complete the first genome-wide survey of RNA exosome substrates in any organism using inducible RNAi lines targeting RRP4 and RRP41 in combination with whole-genome tiling arrays (Chekanova et al., 2007). This pioneering study did notice that, upon exosome knockdown, a group of polyadenylated RNA species referred to as upstream non-coding transcripts (UNTs) exhibited conspicuous accumulation around the 5’-ends of a subset of protein-coding genes. However, the poorly resolved TSS locations available at the time made it unclear whether UNTs generally overlap with 5’-ends of coding transcripts or not. Indeed, in contrast to the situation in mammals in which PROMPTs were also originally found using comparative hybridization to whole-genome tiling arrays and suggested to originate from both forward and reverse strands, the identification of exosome-sensitive UNTs in *Arabidopsis* has not been followed up by more powerful RNA sequencing analyses. Thus, a comparative *Arabidopsis* study using steady-state RNA-seq and RNA 5’-tag sequencing approaches on wild type and mutants defective in nuclear RNA decay components would be a useful complement to nascent RNA studies, not only because of the potential limitations of these techniques, but also because of the original identification of UNTs upon exosome knockdown. Moreover, because of the much higher sensitivity and precision of modern-day sequencing approaches compared to tiling arrays used in the study by (Chekanova et al., 2007), such approaches may also inform on other classes of cryptic transcripts in *A. thaliana,* and on the degradation pathways involved in their rapid turnover.

Although the exosome complex, in association with 3’-5’ exonucleases, is necessary for the decay of many RNA species, its activity *in vivo* depends crucially on adaptor proteins in the DEAD box helicase family that funnel RNA substrates to the exosome (Lykke-Andersen et al., 2009). Plants encode at least three such adaptors: similar to yeast and mammals, SKI2 is required for exosomal degradation of cytoplasmic RNA substrates, including endonucleolytic mRNA cleavage fragments (Branscheid et al., 2015), but in contrast to *S. cerevisiae* and mammals, two different nuclear DEAD-box helicase adaptors are encoded in plant genomes: MTR4 localizes to the nucleolus and has specific functions in ribosomal RNA processing (Lange et al., 2011), while HEN2 localizes to the nucleoplasm and is required for the degradation of a wide range of RNA substrates, including many non-coding RNAs (Lange et al., 2014). In addition to the RNA exosome, less well-characterized nuclear 5’-3’ exonucleolytic pathways exist. These pathways may involve the nuclear 5’-3’ exonucleases XRN2 and XRN3. XRN2 has documented functions in ribosomal RNA cleavage (Zakrzewska-Placzek et al., 2010), while XRN3 has more identified nuclear substrates, including RNA polymerase II-associated RNA and cleavage products by DICER-LIKE enzymes, the former giving rise to defects in transcription termination in *xrn3* mutants (Krzyszton et al., 2018). In addition, the heptameric nuclear Lsm complex, Lsm2-8, a well-established pre-mRNA splicing factor (Bouveret et al., 2000; Tharun et al., 2000), also plays roles in nuclear RNA decapping and, ultimately, decay, presumably via 5’-3’ exonucleolysis (Golisz et al., 2013). In yeast, mammals, and plants, Lsm2-8 shares six of its subunits, from Lsm2 to Lsm7, with the cytoplasmic decapping activator Lsm1-7, such that Lsm8 is the only subunit specific to the nuclear Lsm complex (Perea-Resa et al., 2012).

In this study, we selected a knockout mutant in HEN2, a hypomorphic loss-of-function mutant in the core exosome subunit RRP4, and a knockout mutant in LSM8 for transcriptome-wide studies. We applied transcriptome-wide sequencing of capped 5’ ends of RNAs (CAGE (Takahashi et al., 2012)) and RNA-seq to RNA isolated from mutant and wild type seedlings to identify transcripts subject to preferential nuclear degradation. In agreement with the results of nascent RNA analyses, we also found that the majority of promoters of coding genes only have CAGE signal on the sense strand, even when the exosome complex and its associated helicase system were rendered non-functional. However, clear exceptions to these observations exist: we found 96 mRNA TSSs with strong PROMPT-like configurations that produce exosome-sensitive transcripts, supported by CAGE and RNA-seq, and, in some cases, also by previously obtained GRO-seq data. We also found 113 non-coding regions featuring bidirectional transcription of exosome-sensitive RNAs reminiscent of active enhancers in vertebrates.

While exosome mutants do not feature bidirectional transcription initiation to the same degree as in mammalian genomes, a striking phenotype in *A. thaliana* not present to the same degree in mammalian genomes is the common occurrence of exosome-sensitive RNAs transcribed antisense to genes. Such transcripts commonly initiate in 3’-UTR regions of cognate genes and are ∼1000 nt long, typically unspliced, and tend to occur in genes encoding transcription factors (TFs). In addition, we show that the set of active TSSs in wild type plants presented here has higher accuracy than current TSS annotations such as TAIR10 (Berardini et al., 2015) and, in particular, ARAPORT11 (Cheng et al., 2017). We also demonstrate the prevalence of alternative TSSs in the wild type transcriptome. Our data, therefore, constitute an important resource for gene regulation and RNA degradation research in *Arabidopsis thaliana*.

## RESULTS

### Generation of a comprehensive map of accurate *Arabidopsis thaliana* transcription start sites

Investigations of bidirectional transcription are crucially dependent on highly accurate measurements of TSS locations and activity. As a baseline for further analysis, we purified RNA from 14 day-old *Arabidopsis thaliana* Col-0 wild type seedlings in biological triplicates and prepared CAGE libraries from these RNA samples (Figure 1A). CAGE reads were mapped to the TAIR10 genome assembly with an average of 20.8 M (83.4% of total) uniquely mapped reads per replicate. As described previously, gene TSSs in complex genomes, including *Arabidopsis* (Tokizawa et al., 2017), are often locally dispersed, a phenomenon often referred to as ‘broad promoters’ (Carninci et al., 2006). It is therefore helpful to cluster CAGE tags with closely spaced 5’-ends on the same strand into CAGE tag clusters (TCs). We created tag clusters across all libraries, and calculated the normalized expression of each library as tags per million mapped reads (TPM). Because a TC may be composed of multiple close TSSs, we will use the ‘TC’ term for CAGE initiation events rather than just ‘TSS’, which we will use for annotated 5’-ends and data from other methods. Figure 1B shows an example of CAGE tags identifying the annotated TSS for the Transmembrane protein 97 gene (AT2G32380).

**Figure 1.**
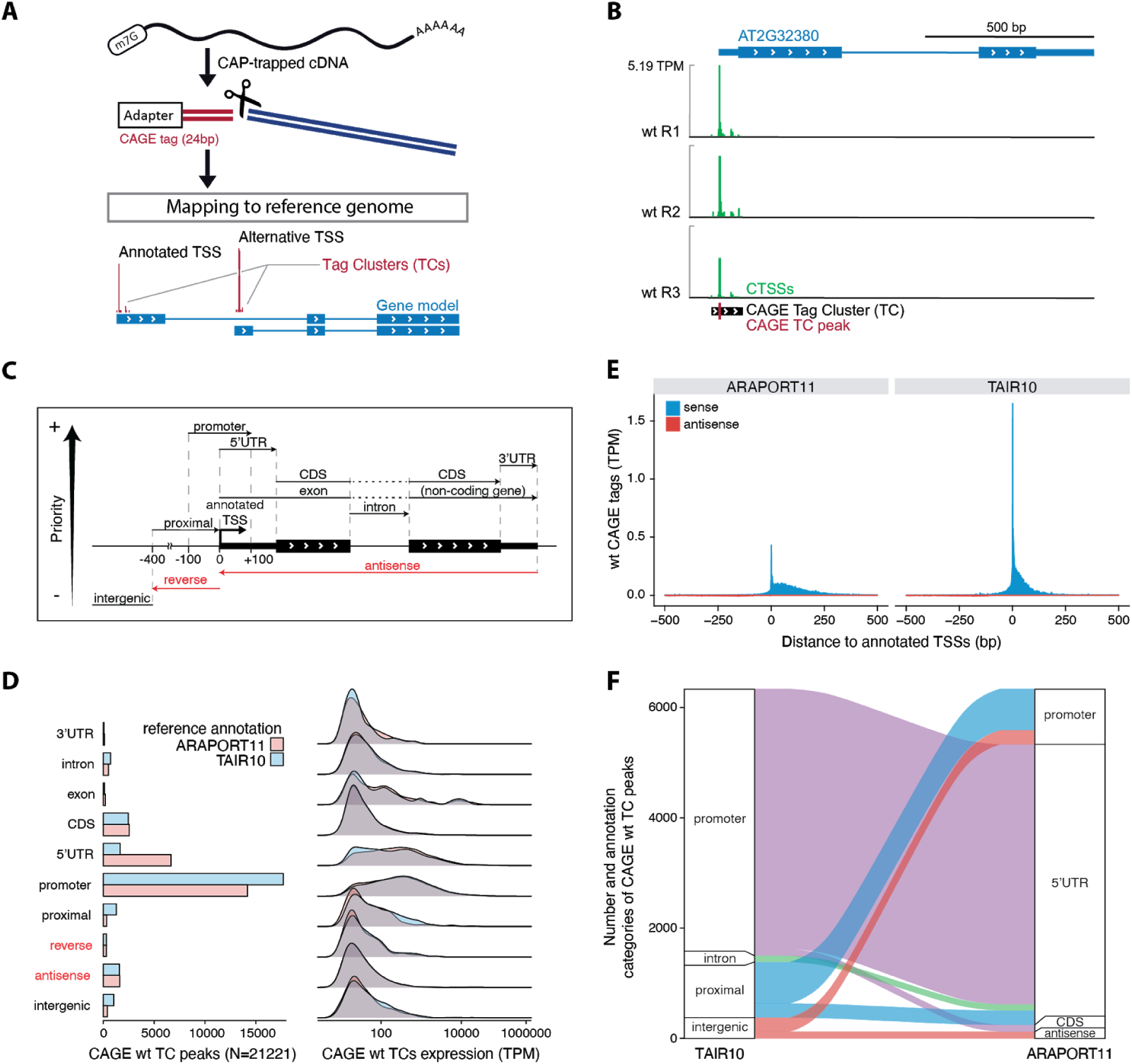
Experimental design and annotation of CAGE TCs. **(A) Conceptual overview of CAGE protocol.** Full-length mRNAs are isolated with cap-trapping and reverse transcribed into cDNA. A CAGE adapter is added to the 5’-end and fragments are cleaved. The remaining CAGE tags are sequenced using a high-throughput platform and mapped to the reference genome. The nearby 5’-ends of CAGE tags are combined into strand-specific tag clusters (TCs). **(B) Example genome browser view of CAGE data.** The AT2G32380 gene model is represented at the top with thin and large blue blocks for untranslated and protein-coding exons, respectively. Chevrons in white show the direction of transcription. Blue lines indicate introns. CAGE tags per million (TPM) signals for each replicate are shown below on the sense strand (green). No antisense CAGE signal was detected for the AT2G32380 locus, so minus strand CAGE signals are not shown. The CAGE TC is shown with a black block, and its peak is indicated with a red tick. **(C) Hierarchical annotation strategy.** Schematic representation of the annotation of CAGE TCs with respect to the reference genome. CAGE TC peaks are used as a proxy for CAGE-detected TSSs. In case a CAGE TC peak overlaps several genomic features, the highest feature in the hierarchy is given priority, as indicated by the arrow (left margin). The promoter region is extended 100 bp in both directions of the annotated TSS, whereas the promoter-proximal region is extended 400 bp upstream (violet). CAGE TCs falling outside of the indicated categories were annotated as intergenic. **(D) Annotation and expression of CAGE TCs.** Left: number of wild type CAGE TCs (X-axis) falling into distinct genomic categories (Y-axis), following the hierarchical annotation strategy defined in C. Right: wild type CAGE expression (X-axis) for the different annotation categories (Y-axis). TAIR10 (blue) and ARAPORT11 (pink) reference annotations were used independently. Annotation categories on the opposite strand of the gene are marked in red. **(E) CAGE footprint at annotated TSSs.** X-axis shows the distance relative to ARAPORT11 (left) and TAIR10 (right) annotated TSSs in bp. Y-axis shows wt CAGE TPM-normalized signal per bp. CAGE sense signal has positive values (blue), while antisense signal has negative values (red). **(F) Differential CAGE TC annotation.** Y-axis shows the number of wt CAGE TCs falling into different annotation categories in TAIR10 vs. ARAPORT11 reference annotation data sets (X-axis). Colors show the link between annotation categories. Only pairs of categories with more than 120 CAGE TCs are shown.

While CAGE mapping is independent of gene annotation, it is often helpful for further analysis to assess into which annotated regions they fall (e.g., to what gene can a given TC and its activity be linked). As in (Thodberg et al., 2018), we devised a hierarchical annotation scheme (Figure 1C) based on gene models from either TAIR10 or ARAPORT11 and counted the number of wild type CAGE TCs located in each type of annotated region (Figure 1D**, left**). As expected, the large majority of TCs fell into annotated promoter regions (+/- 100 bp around the gene model 5’-ends from ARAPORT11 or TAIR10), although ARAPORT11 promoters accounted for a smaller fraction of TCs than TAIR10. All other categories accounted for only a minor fraction of TCs when using TAIR10, but when using ARAPORT11, a much larger fraction of TCs mapped to 5’-UTR regions. As observed previously in human, mouse and fission yeast (Boyd et al., 2018; Bornholdt et al., 2017; Thodberg et al., 2018), CAGE TCs falling into either promoter or 5’-UTR regions had substantially higher expression than other categories (Figure 1D**, right**).

Comparison of the global footprints of CAGE wild type signal anchored at annotated TSSs from ARAPORT11 and TAIR10 showed a wider, downstream-spread distribution of CAGE signal around ARAPORT11 TSSs than for TAIR10, as well as a lower intensity peak (Figure 1E).

In agreement with the initial analyses (Figure 1D, E), the CAGE TCs that were differentially categorized in ARAPORT11 vs. TAIR10 and fell within the annotation category ‘promoter’ in TAIR10 were almost exclusively annotated as 5’-UTR region in ARAPORT11, while CAGE tags falling into the ‘proximal’ region in TAIR10 (−100 to −400 relative to annotated TSS) were frequently annotated as ‘promoter’ in ARAPORT11 (all categories are defined as in Figure 1C, based on respective gene models) (Figure 1F). These observations strongly indicate that TAIR10 TSSs are in general agreement with CAGE TCs, while ARAPORT11 TSSs are, on average, located upstream of CAGE TCs/TAIR10 TSSs.

### Most *Arabidopsis* genes expressed in unchallenged wild type seedlings only use one promoter

The use of alternative promoters or TSSs is an important process for generating RNA isoforms in complex genomes (Davuluri et al., 2008; Valen et al., 2009; Pal et al., 2011). We define alternative TSSs as transcription initiation events that are distant from each other but within the same gene. Therefore, local dispersions of TSSs at a single promoter (so-called ‘broad TSS distributions’ (Carninci et al., 2006)) are not considered alternative TSSs in this analysis. Alternative promoters can generate mRNAs lacking one or more coding exons or upstream open reading frames, and may have an impact on protein abundance and function (Carninci et al., 2006). To assess the extent of alternative promoter usage, we counted the number of wild type TCs overlapping each TAIR10 gene, only retaining those that contributed ≥ 10% of the total CAGE expression across the gene in order to not count minor events. This showed that the vast majority of genes only used one TC (90%, Figure 2A), located in the annotated promoter (Figure 2B), but we also found a substantial number of genes having multiple TCs; for instance, 1632 genes had 2 TCs contributing ≥ 10% of expression (9%, Figure 2A). In cases where at least two TCs were observed, they mostly occurred either in the annotated promoter or in 5’-UTRs (Figure 2B). The few TCs occurring within protein-coding exons were typically not the most highly expressed TCs within the gene. A full list of genes and corresponding coordinates of TCs has been assembled in **Supplementary Data S1,** and two illustrative examples are shown in Figure 2C-D. Figure 2C shows a case in which the most used TC coincides with an annotated shorter protein-coding transcript, while Figure 2D exemplifies a minor TC occurring at the very end of the gene, which is unlikely to yield a functional protein-coding product.

**Figure 2.**
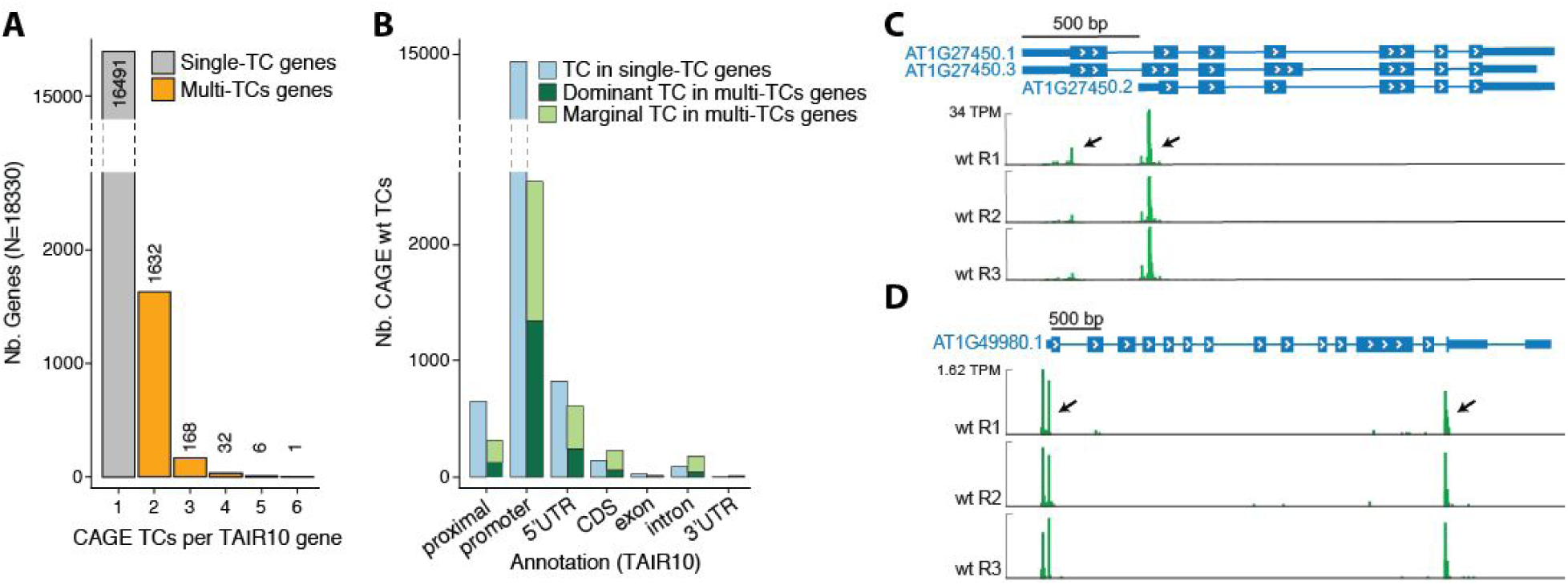
Detection of alternative TSS usage by CAGE. **(A) Alternative TSS occurrence in wild type seedlings.** The X-axis shows the number of sense TCs per TAIR10 gene. For each gene, only TCs accounting for at least 10% of the total number of CAGE tags mapping to the gene are counted. The Y-axis shows the number of genes. Bar colors separate single-TC (grey) from multi-TC genes (orange). The scaling of the Y-axis is split to facilitate visualization. **(B) Location of wild type TCs within genes.** The X-axis shows the classification of sense TCs relative to TAIR10 genes, as in Figure 1C-D. The Y-axis shows the number of TCs by category. Bar colors separate single-TC (blue) from multi-TC (green) genes. Dominant TCs (most expressed within a gene) in multi-TC genes are indicated in dark green, remaining TCs are colored light green. TCs were filtered to contribute at least 10% of the total gene expression. The scaling of the Y-axis is split to facilitate visualization. **(C), (D) Examples of genes with multiple TCs.** Genome-browser images are organised as in Figure 1B, showing the AT1G27450 gene (ATAPT1, ARABIDOPSIS THALIANA ADENINE PHOSPHORIBOSYLTRANSFERASE 1) (C), and the AT1G49980 gene (DNA/RNA polymerases superfamily protein) (D). Tracks show the wild type CAGE data replicates on the same strand, represented as normalized TPM values. Black arrows indicate the CAGE-detected TCs. In C, the most used TC overlaps a known alternative TSS for the gene, corresponding to a shorter transcript lacking one coding exon. In D, the most used TC overlaps the annotated TSS of the gene, whereas the secondary alternative TC is located in the last protein-coding exon, likely producing a non-coding RNA.

This indicates that in unchallenged wild type *Arabidopsis* seedlings, the use of alternative promoters which result in mRNAs encoding different proteins is uncommon. In contrast, several recent studies have documented the widespread occurrence and functional importance of alternative transcription initiation sites in different tissues or upon perception of environmental change, most notably light quality and availability (Yamamoto et al., 2009; Ushijima et al., 2017; Kurihara et al., 2018).

### The TSS annotation in TAIR10 is substantially more accurate than in ARAPORT11

Because we found a discrepancy between ARAPORT11 and TAIR10 TSS annotations, it was important to assess which of these TSS annotations is the most accurate, in particular since our CAGE data were more supportive of the TAIR10 annotation (Figure 1E-F). To do this, we also considered two previous genome-wide TSS data sets: paired-end analysis of transcription sequencing (PEAT-seq) using roots (Morton et al., 2014), and parallel analysis of RNA 5’-ends (nanoPARE) using flower buds (Schon et al., 2018). The majority (62%) of TCs/TSSs was supported by at least two methods, where nanoPARE had the highest fraction of called TCs/TSSs without support from the other methods (25%), compared to CAGE (12%) and PEAT (1%) (Figure 3A, see Methods). The substantial overlap is noteworthy, given that the tissues analyzed differ between studies. Notably, the number of PEAT-seq-defined TSSs was much lower than in the other approaches, and CAGE TCs supported 95% of PEAT-seq TSSs (Figure 3A, B). Further investigation showed that PEAT-seq TSSs had an overall higher expression (as measured by CAGE tags, see Methods) than CAGE TCs. This observation indicates that the two methods detect the same initiation events, but that in addition to the commonly identified set, CAGE also detects TSSs of more lowly expressed genes (Figure 3B).

**Figure 3.**
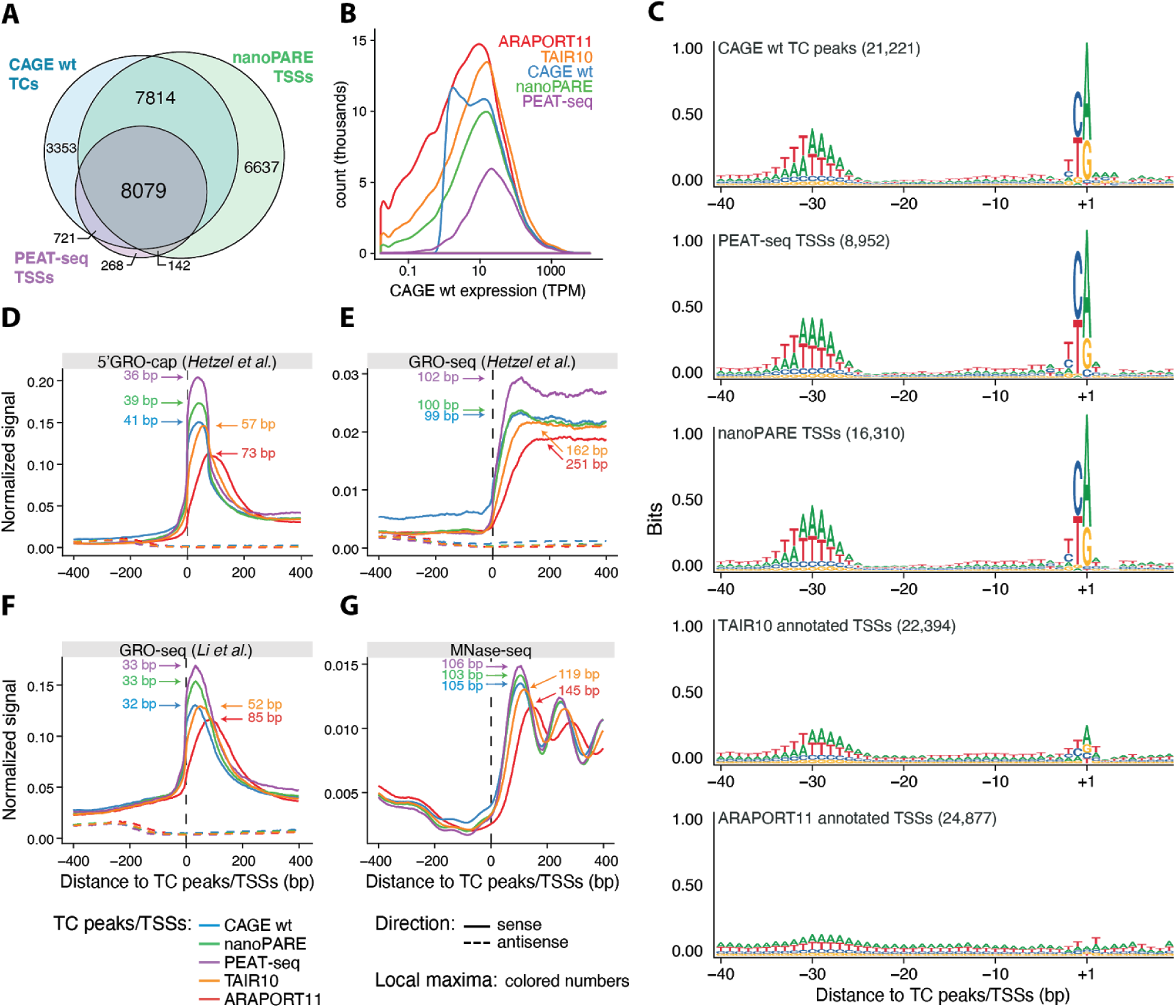
Comparison between TSS data sets. **(A) Comparison of CAGE, PEAT-seq and NanoPARE TCs/TSSs.** Venn diagram showing the number of TCs/TSSs identified by each dataset based on the non-redundant set of all TCs/TSSs (see Methods). **(B) Expression of TCs/TSSs and annotated TSSs across datasets.** The X-axis shows the CAGE expression of TCs/TSSs as TPM (see Methods). The Y-axis indicates the number of TCs/TSSs. Line colors distinguish datasets as indicated. **(C) Sequence patterns around TSSs/TC peaks.** Sequence logos of the genomic sequences surrounding CAGE TC peaks, TSSs defined by PEAT-seq and nanoPARE, and annotated TSSs from TAIR10 or ARAPORT11. The Y-axis shows the information content in bits. The X-axis represents the distance relative to TC peaks/TSSs in bp. **(D), (E), (F) Nascent RNA metaplot around TSSs/TC peaks from respective datasets.** The X-axis shows the position relative to the annotated TSSs from TAIR10 or ARAPORT11, or to TC peaks defined by CAGE, PEAT-seq, and nanoPARE. The Y-axis shows the average normalized signal of 5’ GRO-cap from (Hetzel et al., 2016) (D), and GRO-Seq from (Hetzel et al., 2016) and (Liu et al., 2018) (E, F). Full lines indicate forward strand signal. Dashed lines indicate reverse strand signal. Colors indicate datasets used as anchoring positions (0 at X-axis). Positions of local maxima are indicated with colored text and arrows. **(G) Nucleosome phasing around TSSs/TC peaks from respective sets.** The representation is organized as in D, but in this case, the Y-axis shows the average normalized MNase-seq signal (unstranded data from (Zhang et al., 2015)).

We reasoned that the most accurate TSS annotation set should be best at recapitulating known promoter biology in terms of accessible DNA, sequence characteristics, and data from nascent RNA assays. To make the analyses comparable between sets and focus on TSSs expressed in our samples, we only assessed TCs/TSSs that had a CAGE expression ≥ 1 TPM in at least two replicates in the +/-100 bp region around TCs peaks or annotated TSSs (depending on analysis, see below).

First, we assessed core promoter patterns by constructing sequence logos (Schneider and Stephens, 1990) from DNA sequences around annotated TSSs from TAIR10 and ARAPORT11, TC peaks from CAGE wt, and PEAT-seq and nanoPARE TSS sets (Figure 3C). Sequence logos constructed based on CAGE TCs resulted in the expected sequence patterns observed previously in mammals (Carninci et al., 2006) and later also in *A. thaliana* (Yamamoto et al., 2009; Morton et al., 2014): a strong pyrimidine-purine (PyPu) signal at positions −1/+1 relative to the TC peak, corresponding to the central part of the initiator (Inr) motif, and a TA-rich pattern at positions −35 to −27, corresponding to the TATA box (Figure 3C). TSSs defined by PEAT-seq and nanoPARE showed a highly similar pattern, while TSSs defined by TAIR10-annotated TSSs had weaker TATA-box and pyrimidine-purine dinucleotide patterns. Remarkably, both signatures were almost not visible around ARAPORT11 annotated TSSs, strongly indicating that the TSS annotation of ARAPORT11 poorly captures these central elements of promoter biology.

Second, we assessed the average signal of nascent RNA approaches relative to TCs/TSSs. Three data sets were used: GRO-seq from two different laboratories (Hetzel et al., 2016; Zhu et al., 2018) and 5’ GRO-cap from one (Hetzel et al., 2016)). We observed a consistent shift of GRO-seq and GRO-cap signal maxima in accordance with the above results: TCs/TSSs defined by CAGE, PEAT-seq, and nanoPARE yielded highly similar locations of GRO-seq maxima 35-40 bp downstream of the TSS (Figure 3D-F). For TAIR10-annotated TSSs, a slight 3’-shift in the GRO-cap maximum was seen, and this shift became substantial for ARAPORT11-annotated TSSs (Figure 3D). The same trend was observed using GRO-seq signal from two different laboratories (Figure 3E, F).

Third, Micrococcal Nuclease-seq (MNase-seq) signal (Zhang et al., 2015) from *A. thaliana* seedlings that reports the nucleosome occupancy of genomic DNA showed similar peaks and amplitudes when centered on TSSs defined by CAGE, PEAT-seq, and nanoPARE: In all of these cases, the first peak corresponding to the center of the +1 nucleosome was located on average at 103 to 106 bp downstream of the TSS (Figure 3G). As above, a small 3’-shift was observed when centering on TAIR10 annotated TSSs (+119 bp), and a substantially shifted peak maximum and lower amplitude were observed when using ARAPORT11 annotated TSSs (+145 bp, Figure 3G).

Overall, these results indicate that CAGE, PEAT-seq and nanoPARE are the most accurate TSS sets, where PEAT-seq likely has lower sensitivity, followed by annotated TAIR10 TSSs. In contrast, ARAPORT11 TSSs are misplaced by 128 bp upstream on average, possibly due to the overestimation of transcript lengths, as suggested previously (Schon et al., 2018) . Due to the higher agreement of CAGE, and other 5’-end sets, with TAIR10 annotated TSSs vs. ARAPORT11, we only used TAIR10 annotation in subsequent analyses below.

### Analysis of bidirectional TSS activity at mRNA promoters

Having established an extensive and high-quality set of *Arabidopsis* TSSs, we next focused on the important problem of promoter directionality. To comprehensively map transcripts rapidly degraded by the nuclear exosome, we prepared triplicate CAGE and ribosomal-RNA-depleted total RNA-seq libraries from *hen2-4* and *rrp4-2* mutant seedlings. *hen2-4* is a null allele caused by an exonic insertion of a T-DNA (Lange et al., 2014), while the hypomorphic loss-of-function *rrp4-2* allele encodes an RRP4 protein rendered dysfunctional by the mutation of a conserved residue adjacent to the N-Terminal Domain (NTD), which forms the interface of the core complex with the RRP6 exonuclease (Hématy et al., 2016). Principal component analysis (PCA) on CAGE TC level showed that wild type (as analyzed in Figure 1**-**2), *hen2-4,* and *rrp4-2* samples were separable by the first principal component; *rrp4-2* was the most dissimilar from wild type (Supplementary Figure 1A).

Our first objective was to use the data to assess the occurrence of PROMPT-like, exosome-sensitive transcripts in *Arabidopsis*. In vertebrates, TSSs of protein-coding genes tend to be located at the edges of DNase I hypersensitive sites (DHSs), marking open chromatin regions, close to the +1 and −1 nucleosome. We therefore analyzed the overlap between DHS regions from 14 day-old seedlings (Zhang et al., 2012) and our CAGE data: 89.7% of CAGE tags from wild type fell within DHS regions, which is similar to results from HeLa cells (Andersson et al., 2014b). This motivated the use of our CAGE data together with the DHS-seq data of Zhang et al. (2012).

We first selected DHSs overlapping annotated TSSs of mRNAs. To distinguish sense from antisense CAGE signal, DHS regions were attributed the same strand as the mRNA TSS they overlapped. We then constructed a metaplot of the average CAGE TPM signal using the DNase I maxima (DHS summits, DHSSs) as anchor points. This analysis showed that the sense (mRNA) CAGE signal was on average 25-fold higher than on the antisense signal, regardless of whether nuclear exosomal RNA decay systems were functional (Figure 4A). Specifically, we did not detect a high average enrichment of upstream signal on the reverse strand in exosome mutants, as would be expected if exosome-sensitive PROMPTs were prevalent (Figure 4A). These observations suggest that divergent transcription resulting in exosome-sensitive PROMPTs at promoters of protein-coding genes is not widespread in *A. thaliana*, thus agreeing with (Hetzel et al., 2016) and (Zhu et al., 2018). To ensure that these results were not specific to the CAGE technique, we also plotted RNA-seq reads from wild type and *rrp4-2* mutant seedlings. This produced similar outcomes, albeit with an expected shift downstream of TSSs, due to the inherent characteristics of RNA-seq technique, which tends to miss RNA fragments at the 5’ and 3’ edges (Figure 4B).

**Figure 4.**
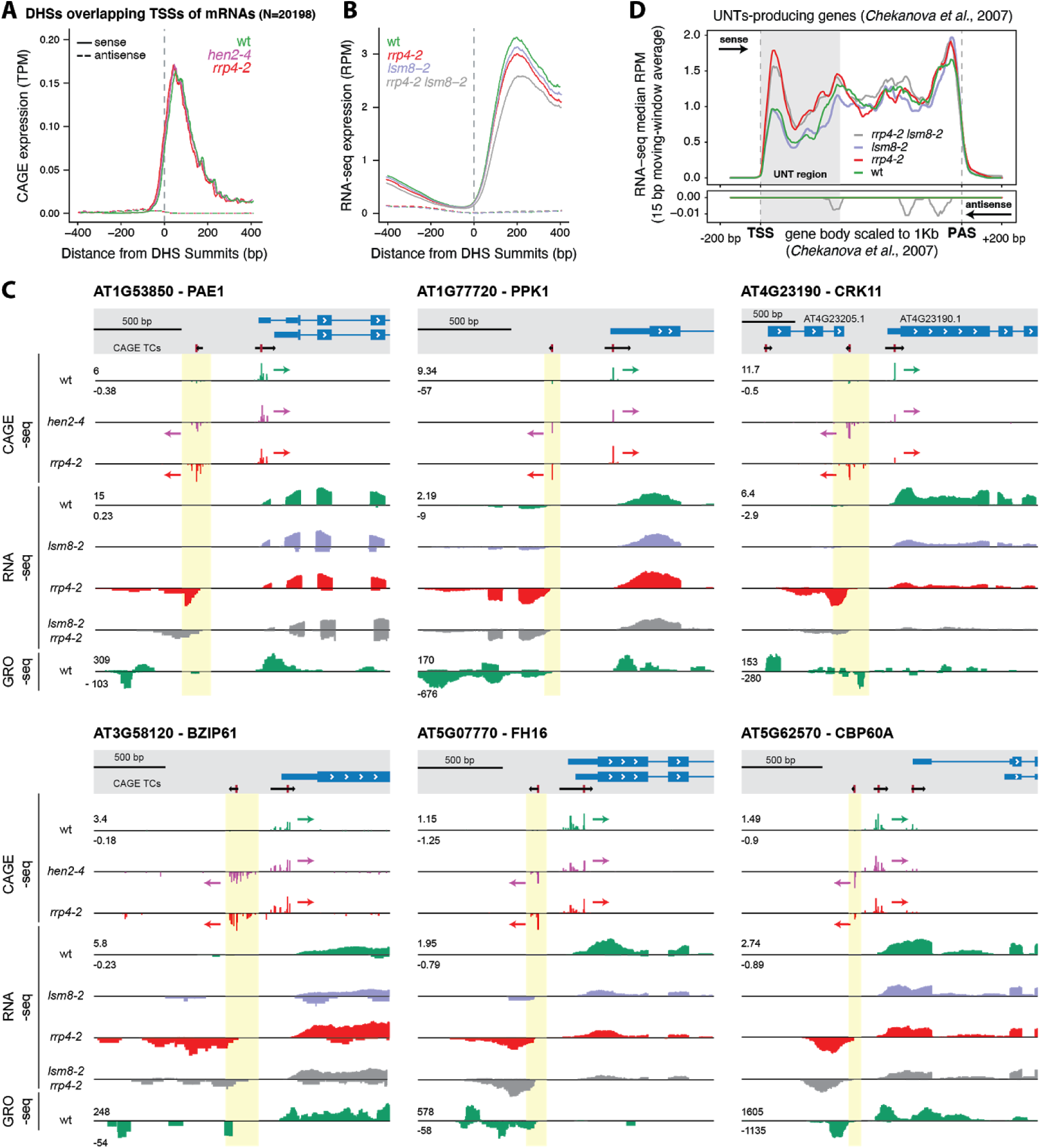
Analysis of bidirectional transcription initiation at promoters of *Arabidopsis* protein-coding genes. **(A) CAGE signal at DHSs overlapping annotated TSSs of protein-coding genes.** The Y-axis shows the average normalized CAGE signal in TPM, smoothed with a 15 bp moving window. The X-axis represents the distance (in bp) relative to the maxima of the DHS peaks that overlap TAIR10 annotated TSSs. Colors distinguish genotypes as indicated. Full and dashed lines indicate sense and antisense strands, respectively. **(B) RNA-seq signal at DHSs overlapping annotated TSSs of protein-coding genes.** Organized as A: Y-axis shows normalized average RNA-seq signal as reads per million mapped reads (RPM). **(C) Examples of bidirectional transcription at promoters of protein-coding genes.** Six cases of bidirectionally transcribed promoters of protein-coding genes organized as in Figure 1B are shown. Top: TAIR10 transcript models (blue) and CAGE TCs (black; TC peaks in red). Tracks show normalized CAGE, RNA-seq, and GRO-seq (from (Liu et al., 2018)) signals. Colors differentiate genotypes. Signals on forward strand have positive values, signals on reverse strand have negative values. Arrows show the direction of transcription for CAGE TCs. PROMPT TSS regions are highlighted in yellow. **(D) RNA-seq metaplot for UNT-producing genes.** The Y-axis shows the median sense and antisense normalized RNA-seq signal (10 bp moving average), colored by genotype. Positive values denote sense signal, and negative values denote antisense signal (also indicated by arrows). X-axis shows scaled position in UNT-producing genes, where gene bodies were scaled to 1 kb. Vertical dashed lines indicate TAIR10 annotated TSS and polyadenylation site (PAS), flanked by 200 unscaled base pairs (bp). Grey box indicates the UNT region.

### Inactivation of nuclear decapping does not promote detection of PROMPTs

Two different scenarios may explain the overall low CAGE and RNA-seq signal in exosome mutants upstream of mRNA TSSs on the reverse strand: (i) there is on average little transcription initiation in these regions (in agreement with the lack of GRO-seq signal (Hetzel et al., 2016)), or (ii) transcription of PROMPTs may occur, but *A. thaliana* may employ redundant RNA decay systems for their degradation, precluding their detection by simple inactivation of the nuclear exosome. To explore the latter possibility, we extended the RNA-seq experiments to also profile seedlings defective in nuclear decapping (and hence 5’-3’ exonucleolysis), or in both nuclear decapping and exosome activities. To this end, we used the *lsm8-2* knockout mutant (Perea-Resa et al., 2012; Golisz et al., 2013) and also constructed an *rrp4-2 lsm8-2* double mutant in which both RRP4 and LSM8 were defective. RNA-seq metaplot of both *lsm8-2* single and *rrp4-2 lsm8-2* double mutants were highly similar to the wild type (Figure 4B), strongly indicating that LSM8 is not part of a redundant pathway for PROMPT degradation.

### Clear PROMPTs can be detected at individual genes

Despite the low average signal upstream of genes on the reverse strand in metaplots, visual genome-browser inspection allowed us to detect cases of exosome-sensitive transcripts with the characteristics of mammalian PROMPTs. To identify such cases systematically, we analyzed the same TSS-overlapping DHSs as above and required *rrp4-2* CAGE expression on the reverse strand upstream of mRNA TSSs to be two-fold higher than that of wild type, and statistically significant (log_2_ fold change □ 1, *FDR* □ 0.05). Despite the conservative selection criteria, this resulted in 96 unique regions (see examples in Figure 4C), (**Supplementary Data S1**). We next asked to what degree previously published nascent transcription data supported these clear PROMPT examples, and defined PROMPT regions with evidence of transcription as regions with 2-fold more signal than the genome-wide average (see Methods). Using this criterion, about half of the identified PROMPTs were supported by nascent transcription data (on average 52%, depending on the nascent RNA dataset, see Supplementary Figure 1B) (Hetzel et al., 2016; Liu et al., 2018). Given the high thresholds we have employed for CAGE data, these results indicate that many PROMPTs do indeed escape detection by the available nascent transcriptome data.

Taken together, our data indicates that most TSSs of protein-coding genes appear to be unidirectionally transcribed, in agreement with previous conclusions. However, clear exceptions exist, where highly exosome sensitive transcripts are initiated at locations similar to that of the much more widespread PROMPTs in vertebrates. It is therefore possible that initiation of transcription by RNA polymerase II is fundamentally similar in plants and vertebrates, but that plants have evolved mechanisms in addition to selective nuclear RNA decay to ensure a directional promoter output, a point that we elaborate on in the Discussion.

### Properties of PROMPTs and their associated genes

We next sought answers to three questions to better characterize the identified PROMPT-producing regions. First, since the set of PROMPT-associated genes was small, we asked whether PROMPT production could be linked to a specific biological function of the associated coding genes. Gene ontology (GO) term analysis showed no significant over-representation of GO terms, suggesting that there is no apparent functional link amongst PROMPT-associated genes. Second, since enrichment of poly(A) sites and depletion of 5’-splice sites is pronounced in mammalian PROMPT regions compared to mRNAs (Ntini et al., 2013; Almada et al., 2013), we asked whether the same patterns were observed in the identified *Arabidopsis* PROMPT regions.

In contrast to mammals, we only observed a small enrichment of predicted poly(A) sites (AWTAA consensus) after 200 bp in PROMPT regions and no difference in the occurrence of predicted 5’-splice sites (see Methods) (Supplementary Figure 1C). By contrast, we did observe a significant enrichment of predicted 3’-splice sites (Kolmogorov-Smirnoff test, D = 0.33, *P* = 0.0009) (Supplementary Figure 1C), although the underlying mechanistic cause of this enrichment remains unclear. The modest enrichment in poly(A) sites may in part be due to the fact that plants use more diverse, and less well characterized, poly(A) sites than mammals (Li and Du, 2014). Third, we asked whether there was any relation between PROMPT-producing regions and the UNT-producing genes identified by (Chekanova et al., 2007). None of the 96 genes with exosome-sensitive PROMPT-like production belonged to the set of UNT-producing genes. Thus, antisense PROMPT production and sense UNT production are not linked. We note, nonetheless, that the UNTs were clearly detected in our experiments as regions with higher RNA-seq signal in *rrp4-2* downstream of TSSs than in wild type (Figure 4D).

### Evidence for bidirectional TSS activity at intronic and intergenic space

We next used the data to examine possible bidirectional transcription of intronic or intergenic loci, as this has been shown in vertebrates and insects to be a powerful predictor of enhancer regions and enhancer activity (Andersson et al., 2014a; Rennie et al., 2018). Visual inspection also showed several such cases in *Arabidopsis* (examples in Figure 5A). To identify such regions systematically, we used the same approach as in (Thodberg and Sandelin, 2019; Thodberg et al., 2019) to locate genomic regions with a balanced bidirectional CAGE signal. A total of 113 bidirectional clusters was identified, of which 78 were intronic, and 35 were intergenic (**Supplementary Data S1**). On average, such regions were more highly expressed in *rrp4-2* or *hen2-4* compared to wild type, and exhibited clear average bidirectional CAGE transcription in all mutants except *lsm8-2*, both in CAGE and RNA-seq data (Figure 5B). A similar but less distinct trend could be observed in 5’ GRO-cap data (Figure 5C). RNA polymerase II (RNAPII) ChIP-seq signal from 12 day-old seedlings (Cortijo et al., 2017) was also enriched at the edges of NDRs; in intronic regions, these were highly balanced while there was a surprising minus-strand bias in intergenic regions (Figure 5D).

**Figure 5.**
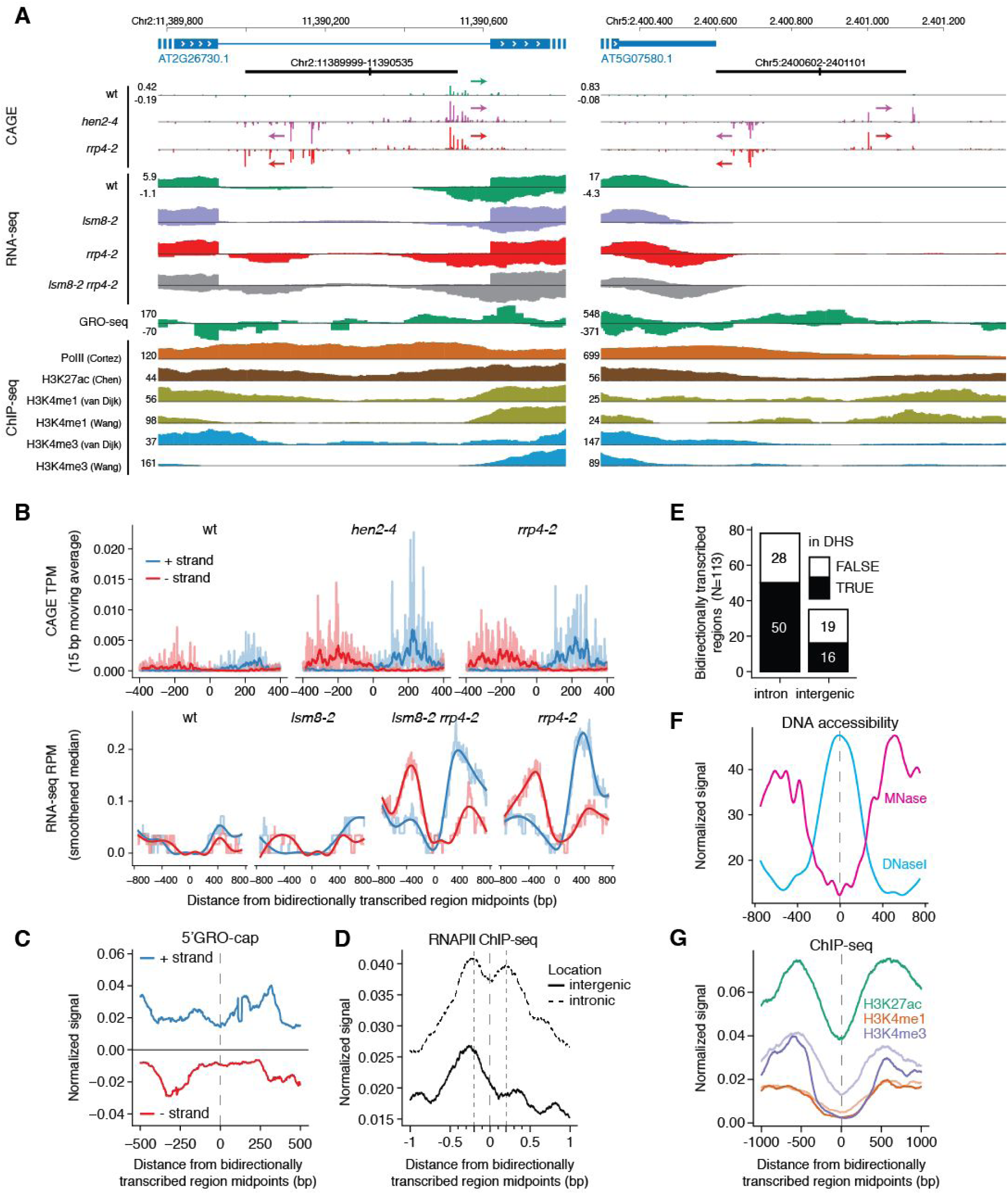
Evidence of bidirectionally transcribed intronic and intergenic regions in *Arabidopsis*. **(A) Examples of bidirectionally transcribed regions.** Genome browser views organized as in Figure 4D. Examples of intronic (left) and intergenic (right) bidirectionally transcribed regions are shown with the TAIR10 gene annotations (blue, top). Tracks from top to bottom: CAGE-seq, RNA-seq, 5’ GRO-cap (Hetzel et al., 2016), and ChIP-seq normalized signals. Positive signals indicate sense transcription; negative signals indicate antisense transcription. Arrows indicate CAGE TCs. **(B) CAGE and RNA-seq at bidirectionally transcribed regions.** The Y-axis shows CAGE (top) and RNA-seq (bottom) signal, colored by strand. Light color indicates average normalized signal. CAGE dark color indicates 15 bp moving-window average. RNA-seq dark color indicates a spline-smoothed signal. The X-axis shows the distance relative to the midpoint position of 113 bidirectionally transcribed regions, in bp. Columns separate the different genotypes, as indicated. **(C) 5’ GRO-cap at bidirectionally transcribed regions.** The X-axis is organized as in B. The Y-axis shows the average 5’ GRO-cap normalized signal from (Hetzel et al., 2016). Blue color and positive values indicate plus strand. Red color and negative values indicate minus strand. **(D) RNAPII at bidirectionally transcribed regions.** The X-axis is organized as in B. The Y-axis shows the average normalized RNA Pol II ChIP-seq signal from (Cortijo et al., 2017). Full and dashed lines indicate bidirectionally transcribed regions annotated as intergenic or intronic, respectively. Vertical dashed lines indicate RNAPII maxima. **(E) Overlap of bidirectionally transcribed regions with DHSs.** The X-axis shows the annotation of bidirectionally transcribed regions. Y-axis counts the number of bidirectionally transcribed regions. Colors indicate whether the bidirectionally transcribed regions overlap with a DHS. **(F) DNA accessibility at bidirectionally transcribed regions.** The X-axis is organized as in B. The Y-axis shows the average normalized MNase-seq and DNAse-seq signal from (Zhang et al., 2016), separated by color. MNase-seq signal is proportional to the nucleosome occupancy, whereas DNase-seq signal denotes NDRs. **(G) Histone marks at bidirectionally transcribed regions.** The X-axis is organized as in B. Y-axis: normalized signal of H3K27ac from 12 day-old seedlings (Chen et al., 2017) (green), H3K4me1 from 4 week-old leaves (orange), and H3K4me3 from 3 week-old plants (violet). Color transparency distinguishes same histone marks but originating from distinct authors (light color: (van Dijk et al., 2010); dark color: (Wang et al., 2015)).

To further characterize these bidirectionally transcribed loci, we assessed their chromatin state using published data: the majority of bidirectional CAGE sites overlapped with a DHS (Figure 5E) and were centered on accessible DNA, as measured by DNase and MNase-seq (Figure 5F). Histones adjacent to the NDRs were enriched for H3K27ac (Chen et al., 2017) and to a lesser degree, H3K4me1 and H3K4me3 (Wang et al., 2015; van Dijk et al., 2010) (Figure 5G). In vertebrates, high levels of H3K4me1 are common in enhancer regions, but this histone modification has been associated with active transcription elongation in *Arabidopsis* (Nielsen et al., 2019). Overall, our analyses indicate that some DNase I hypersensitive intergenic and intronic loci in *Arabidopsis* feature bidirectional transcription initiation at NDR edges. The majority of these loci produce exosome-sensitive RNAs and have enhancer-associated chromatin marks. While such patterns are predictive of enhancer activity in vertebrates, their possible enhancer activity remains to be tested in *Arabidopsis*.

### Identification of sets of mRNAs sensitive to *rrp4-2* and *hen2-4*

We reasoned that while PROMPTs and eRNAs are rarely detected, even when impairing two distinct nuclear RNA degradation systems, a systematic comparison of CAGE and RNA-seq data from exosome mutants and wild type may identify hitherto unknown transcripts in other regions of the genome. Because we wanted to find exosome substrates, we focused on transcripts that were more abundant in either mutant. We first identified CAGE TCs whose expression was significantly higher in *hen2-4* or *rrp4-2* compared to wild type (log_2_ fold change □ 1, FDR □ 0.05, by the limma method (Ritchie et al., 2015), see **Supplementary Table S2**).

This resulted in a total of 1747 up-regulated TCs in both mutants combined. All up-regulated CAGE TCs showed the expected nucleotide distribution around TSSs with enriched TATA and pyrimidine-purine dinucleotide patterns at −30 and +1, respectively (Supplementary Figure 2A), making it likely that they represent genuine TSSs. TCs up-regulated in *hen2-4* were less numerous, and in 92% of the cases, they were also significantly changed in *rrp4-2* compared to wild type (Fisher’s exact test, *P* = 0) (Figure 6A). The few TCs (N=73) significantly up-regulated only in *hen2-4,* but not *rrp4-2* (Figure 6B) may represent transcripts whose exosome targeting requires HEN2, yet their exosomal decay still works in the hypomorphic *rrp4-2* mutant. Alternatively, HEN2 may have as-yet undescribed functions independent of the exosome. This latter scenario is supported by the fact that only 2 of the 73 transcripts were found in the set of 848 transcripts up-regulated upon inducible RNAi knockdown of RRP4 (Chekanova et al., 2007).

**Figure 6.**
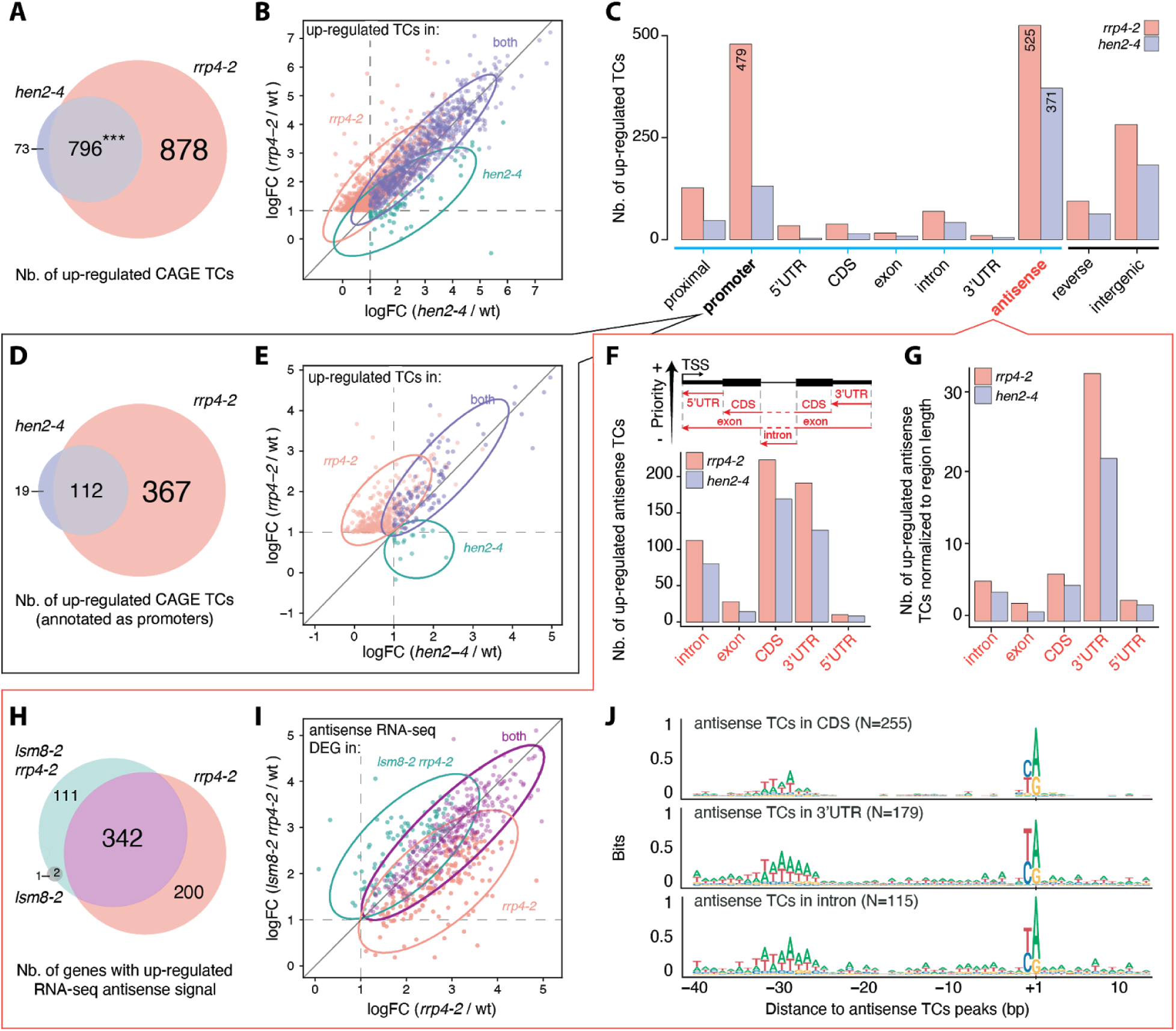
Systematic characterization of exosome-sensitive transcripts in *Arabidopsis*. **(A) Overlap of TCs up-regulated in *rrp4-2* and *hen2-4*.** A CAGE TC was defined as up-regulated if differentially expressed in mutant *vs.* wild type (log_2_ fold change □ 1, *FDR* □ 0.05). **(B) Fold-change relation of up-regulated TCs across mutants.** X-and y-axes show log_22_ fold change of *hen2-4* and *rrp4-2* mutants *vs.* wild type, respectively. Each dot represents an up-regulated CAGE TC, colored by whether its up-regulation was specific or shared between mutants. Ellipses group the majority of TCs in each class. **(C) Annotation of CAGE TCs up-regulated in *rrp4-2* and *hen2-4*.** X-axis: number of TCs overlapping each TAIR10 annotation category as bars, colored by mutant. Y-axis shows gene annotation categories, as in Figure 1D. ‘Antisense’ (red label) refers to any position on the opposite strand of an annotated gene. Remaining figures constitute analyses of subsets of this figure: TCs falling into promoter regions (D, E) and antisense to genes (F-J), as indicated by black and red callouts **(D) Overlap of mRNA promoter TCs up-regulated in *rrp4-2* and *hen2-4.*** Organized as in panel A, but only assessing TCs falling into annotated mRNA promoter regions. **(E) Fold-change relation of up-regulated mRNA promoter TCs across mutants.** Organized as in panel B, but only assessing TCs falling into annotated mRNA promoter regions. **(F) Annotation of antisense TCs up-regulated in *rrp4-2* and *hen2-4***. Organized as in C, but using the antisense annotation hierarchy as shown above the bar plot, where strandedness of TAIR10 features is inverted to allow annotation of antisense TC locations. **(G) Enrichment of antisense up-regulated TCs by annotation category.** Organized as in F, where X-axis indicate the number of *rrp4-2* and *hen2-4* up-regulated antisense TCs normalized to the genome-wide length of the respective regions (Mbp). **(H) Overlap between genic antisense transcripts up-regulated in *rrp4-2*, *lsm8-2,* and *rrp4-2 lsm8-2*.** The RNA-seq signal, antisense to annotated genes, was used to identify transcripts up-regulated in a mutant compared to wild type. The same differential expression cutoffs as in A were used. **(I) Fold-change relation between antisense up-regulated genes across mutants.** Organized as in B, but using RNA-seq data antisense to a TAIR10 annotated gene. X axis show *rrp4-2* mutants *vs.* wild type fold change, Y-axis shows the double mutant *rrp4-2 lsm8-2* fold change, both in log scale. **(J) Sequence patterns of up-regulated antisense TCs.** Organized as in Figure 3C. Logos show the sequence properties around antisense CAGE TC peaks, divided by their annotation. Only categories with more than 100 exosome-sensitive antisense TC peaks are shown.

We next classified all TCs up-regulated in either *rrp4-2* or *hen2-4* by their overlap with the TAIR10 reference annotation (as in Figure 1C) (Figure 6C). This produced two surprising outcomes.

First, a large number (N=479) of TCs corresponding to sense mRNA TSSs (promoters) were up-regulated in the *rrp4-2* mutant. Some of these TCs (N=112) were also up-regulated in *hen2-4*, but the majority (N=367) was only up-regulated in *rrp4-2,* while an additional 19 TCs were only upregulated in *hen2-4* (Figure 6D, E). The up-regulation could either be a consequence of transcriptional induction as an indirect effect of the *rrp4-2* mutation or be more directly related to reduced exosome activity if the decay of this group of mRNAs involves a substantial contribution from the exosome. In the first scenario of transcriptional up-regulation, one might expect the mRNAs to encode functionally related proteins, but we found no significant enrichment of GO terms in the group of genes associated with the up-regulated TCs. A more direct involvement of the exosome in the decay of these two groups of mRNAs would be of interest: The group of 131 mRNAs up-regulated in *hen2-4* may be appreciably affected by nuclear mRNA quality control, akin to the Poly(A)-tail exosome targeting (PAXT) pathway described in mammals (Meola et al., 2016), while the up-regulation of the larger group of 367 mRNAs specifically in *rrp4-2* may be explained by a substantial contribution of the exosome to their cytoplasmic decay. Notably, the average expression of all of these moderately exosome-sensitive TCs of mRNAs was higher than most other categories, perhaps suggesting that the decay of highly expressed mRNAs tends to implicate the exosome (Supplementary Figure 2B).

The second surprising outcome of our analysis of exosome-sensitive TCs was related to a group of 525 CAGE TCs that were up-regulated in *rrp4-2* vs. wild type. The majority of these TCs (N=347) were also up-regulated in *hen2-4*. These TCs were neither PROMPTs nor bidirectionally transcribed intergenic and intronic loci analyzed above, but were located on the reverse strand within gene bodies, thus representing antisense RNAs (Figure 6C, red highlight). To annotate these antisense TCs more accurately, we used the same hierarchical procedure as above, but with inverted annotation strand. This analysis showed that the exosome-sensitive antisense TCs tended to overlap coding exons (CDS) and 3’-UTRs of mRNAs (Figure 6F). Normalization of the number of TCs to the average lengths of CDSs and 3’-UTRs showed that the 3’-UTR category was most enriched for up-regulated antisense CAGE TCs (Figure 6G). More generally, antisense TCs had similar average expression and fold change across annotation categories, which in turn were generally similar to that of other exosome-sensitive TCs (Supplementary Figure 2C).

To verify the existence of such antisense RNAs, we analyzed RNA-seq reads within but antisense to genes, comparing mutants with the wild type. This analysis yielded higher numbers of differentially expressed transcripts than the CAGE-based analysis: 656 genes were up-regulated (log_2_ fold change □ 1, *FDR* □ 0.05, see Methods) in either *rrp4-2, lsm8-2* or *rrp4-2 lsm8-2* mutants *vs.* wt on the antisense strand (**Supplementary Data S2**). Only three transcripts were up-regulated in the *lsm8-2* single mutant. For the remaining transcripts, 52% were up-regulated in both *rrp4-2* and *rrp4-2 lsm8-2* mutants, while relatively large sets were significantly upregulated only in *rrp4-2* single mutant (30%) or the *rrp4-2 lsm8-2* double mutant (17%) (Figure 6H). Plotting the log_2_ fold-change of *rrp4-2 vs.* wt against the log_2_ fold-change of *rrp4-2 lsm8-2* vs. wt revealed that genes found to be differentially expressed in the *rrp4-2 lsm8-2* mutant had the same trend in *rrp4-2* single mutants (Figure 6I). Thus, we found no clear cases of redundancy between LSM8-dependent and exosome-dependent decay pathways. The antisense CAGE TCs may represent genuine initiation events or partially degraded, recapped RNAs (Affymetrix ENCODE Transcriptome Project and Cold Spring Harbor Laboratory ENCODE Transcriptome Project, 2009). We reasoned that if most TCs are genuine TSSs, they should have a similar distribution of core promoter elements as other TCs. Indeed, the great majority of antisense TCs, regardless of overlap category, displayed a strong Inr as well as TATA motif (Figure 6J), suggesting that the exosome-sensitive, antisense TCs represent *bona fide* TSSs.

### Characterization of exosome-sensitive antisense RNAs initiating towards 3’-ends of genes

Antisense RNAs initiating within 3’-regions of genes encoding developmental regulators have been shown to play essential roles in developmental transitions in plants, including germination and flowering (e.g. (Fedak et al., 2016)). Given this, and the unexpected prevalence of exosome-sensitive TSSs in 3’-regions of genes, we decided to characterize these TSSs and their cognate RNAs in more detail. A total of 354 CAGE TCs antisense to an annotated 3’-UTR region were found across samples using the same expression cutoffs as above, of which 197 (56%) were up-regulated in at least one of the mutants compared to wt (Figure 6D). Interestingly, genes with an up-regulated CAGE TC antisense to their 3’-UTR showed a weakly significant enrichment in ‘*sequence-specific DNA binding’* GO term (GO:0043565, *FDR* = 0.02, 8.7% of genes), indicating that genes encoding transcription factors are more prone to have antisense transcription initiation within 3’-UTRs. Examples of exosome-sensitive antisense transcripts and TCs are shown in Figure 7A.

**Figure 7.**
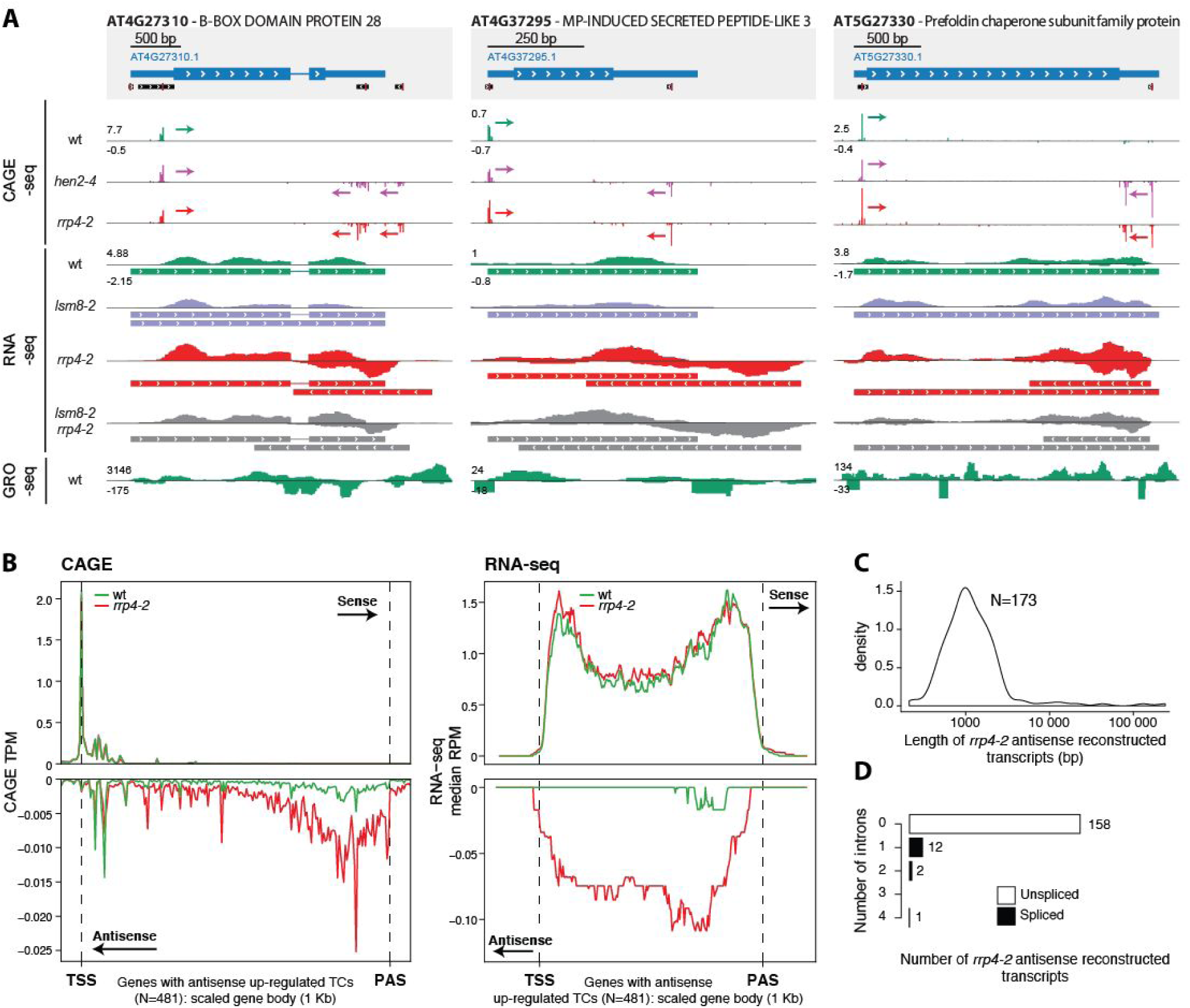
Characterization of exosome-sensitive 3’-UTR antisense TCs. **(A) Examples of exosome-sensitive 3’-UTR antisense TCs**. Genome-browser views are arranged as in Figure 3C. *De novo* assembled antisense transcripts using RNA-seq data sets are also shown for the mutants in which the transcripts are detectable. **(B) Metaplots of CAGE and RNA-seq signal at genes with up-regulated antisense TCs.** The Y-axis shows the average or median normalized read count in 5 bp windows across 481 genes with an antisense, exosome-sensitive TCs. Gene bodies are scaled to 1 kb (x-axes). Colors distinguish wild type from the *rrp4-2* mutant. Reads on sense strand have positive y-values, reads on antisense strand have negative y-values. From left to right, dotted vertical lines indicate the annotated transcription start site (TSS) and poly(A) site (PAS). **(C) Length distribution of 173 *de novo* reconstructed transcripts, antisense to a gene, and up-regulated in the *rrp4-2* mutant.** The X-axis shows length in bp, log-scaled. Y-axis shows distribution density. **(D) Number of detected introns in exosome-sensitive *de novo* reconstructed antisense transcripts for the *rrp4-2* mutant.** The X-axis indicates the number of exosome-sensitive *de novo* assembled antisense transcripts for the *rrp4-2* mutant. The Y-axis shows the number of introns detected in these transcripts.

The CAGE TCs antisense to 3’-UTRs showed no specific bias in terms of location within the 3’-UTR, but were substantially more likely to occur in longer 3’-UTRs (Supplementary Figure 3A, B). Since the *Arabidopsis* genome is compact, we hypothesized that the exosome-sensitive antisense TCs might be PROMPTs from closely located downstream genes. However, this phenomenon appears to be rare: Only 16 instances were identified in which the maximal distance between a 3’-UTR antisense TC and its closest downstream mRNA TSS was 300 bp, and this number only increased to 39 instances when a maximal distance of 500 bp was allowed (Supplementary Figure 3C). More generally, there was no relation between the intergenic distance between genes (3’ end of gene to next 5’ or 3’ gene end downstream) and whether the 3’-UTRs had exosome-sensitive antisense CAGE TCs (Supplementary for Figure 3D).

Because we had both CAGE and RNA-seq data, we analyzed RNA-seq read density across sense and antisense strands for the 481 genes having at least one differentially up-regulated antisense CAGE TC in *rrp4-2* or *hen2-4* (the set is larger than above, because antisense CAGE TCs within other regions than 3’-UTRs were included). This showed that in the exosome mutants, the antisense RNA-seq reads were uniformly distributed across the gene body except 5’-and 3’-ends, but the antisense signal was on average 14.5-fold lower than that of the sense strand (Figure 7B, right). Conversely, CAGE antisense reads resided in the last *∼* 20% of the gene (Figure 7B, left), consistent with their dominant 3’-UTR overlap observed above. Lastly, we leveraged the RNA-seq data to investigate the length and splicing status of *rrp4-2* up-regulated antisense transcripts. To assess this in as unbiased way as possible, we used strand-aware *de novo* transcript assembly for RNA-seq reads (see Methods and **Supplementary Data S3**) in *rrp4-2*, and then plotted the length of novel antisense transcripts (Figure 7C) and the number of introns they included (Figure 7D). The length of antisense transcripts had a large variance, but the most common length was ∼1000 bp, and the vast majority (91.3%) was unspliced. An important caveat with this analysis is that *de novo* transcript assembly is only possible if enough RNA-seq reads are present. This means that these properties are measured only through more highly expressed transcripts, and length estimates may be conservative. Taken together, our analyses revealed that a substantial number of genes in *Arabidopsis thaliana* features antisense transcription of unspliced, exosome-sensitive antisense RNAs, a property that is more common in transcription factor genes.

### The *DOG1/asDOG1* locus exemplifies a complex TSS organization with a functional antisense lncRNA

As a final example of the utility of our data, we analyzed the *DELAY OF GERMINATION1* (*DOG1*) locus in detail. The DOG1 protein is an important developmental regulator that controls seed dormancy such that *dog1* loss-of-function mutants exhibit uncontrolled seed germination, even during fruit development, while DOG1 over-expressers are unable to break dormancy (Fedak et al., 2016). Crucially, DOG1 expression is regulated in *cis* by a lncRNA, asDOG1, which initiates at an alternative 3’-UTR in exon 2 (Fedak et al., 2016). Overlaying our RNA-seq and CAGE data shows the overall complexity and exosome effects at this locus (Figure 8). First, CAGE data shows that DOG1 has two alternative sense strand TSSs: the first corresponds roughly to TAIR10 annotation, while the second is located within exon2, producing an uncharacterized transcript. Interestingly, both CAGE and RNA-seq shows that these transcripts have a degree of exosome sensitivity, which is uncommon for mRNAs. Second, we identify an antisense CAGE TC very close to the previously annotated TSS of asDOG1: both CAGE and RNA-seq data shows that asDOG1 is highly exosome-sensitive (notably, TAIR10 annotation suggests that this TSS lies within the coding region of exon 2, while (Fedak et al., 2016) showed that this region has a dual function as a 3’-UTR for a shorter mRNA isoform). Consistent with our results above, *de novo* RNA-seq isoform reconstruction showed that asDOG1 is not spliced and extends to the annotated TSS of DOG1 (estimated length of 1223 bp), agreeing with the RACE experiments from the original study. The case study of the DOG1/asDOG1 locus shows that the well-established *cis*-regulator of gene expression asDOG1 is one example of the set of exosome-sensitive antisense RNAs that we identified here. Therefore, it is plausible that more of these lncRNAs, or the act of their transcription, have regulatory potential. This hypothesis is particularly appealing because of the enrichment of genes encoding transcription factors in the group of genes giving rise to exosome-sensitive antisense transcripts.

**Figure 8.**
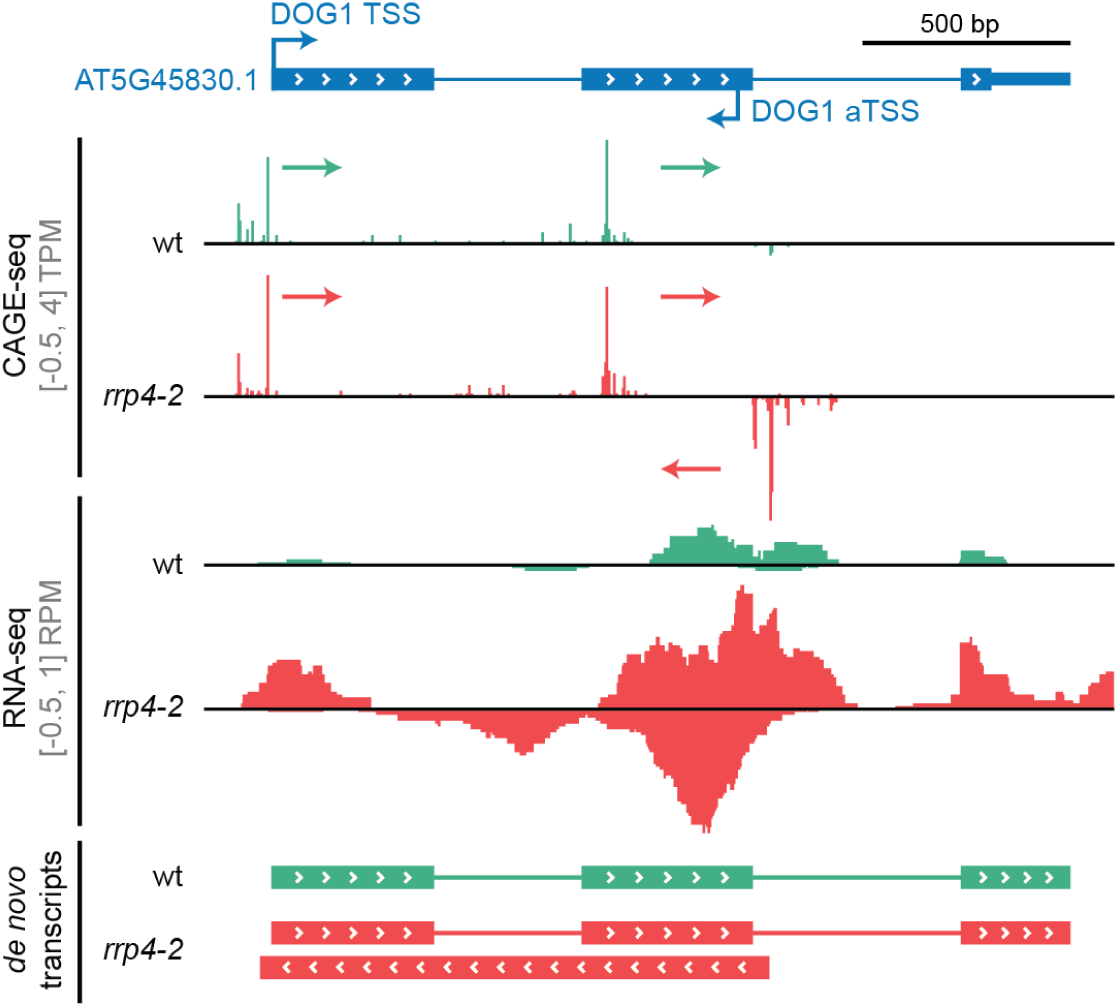
Example of a complex antisense region with a functional antisense lncRNA: the *DOG1/asDOG1* locus. Genome-browser view organised as in Figure 4C. The AT5G45830 (*DOG1*, *DELAY OF GERMINATION1*) gene model is represented at the top with arrows marking the sense annotated TSS and an antisense TSS (aTSS) in the second exon from (Fedak et al., 2016). Tracks show normalized CAGE and RNA-seq signal, and *de novo* reconstructed transcripts using respective RNA-seq data sets. Signals and transcripts are colored by genotype. CAGE-detected TCs and their transcriptional direction are indicated with colored arrows.

## DISCUSSION

### Divergent transcription at genes is uncommon, but not absent in Arabidopsis

In this study, our initial aim was to characterize the extent of bidirectional transcription at promoters of protein-coding genes as well as in intergenic and intronic regions in *Arabidopsis*, given their frequent occurrence in vertebrates. We did this by using two complementary steady-state RNA methods (CAGE and RNA-seq), where critical nuclear degradation enzymes were rendered non-functional. Our results mostly agree with previous efforts using nascent RNA methods (Hetzel et al., 2016): PROMPTs are rare in *Arabidopsis*, but not non-existent: using conservative thresholds, we identified nearly 100 PROMPTs, which are clear exosome substrates. Notably, such cases were not reported previously in studies using nascent RNA methods, even around half of them had nascent RNA support.

The reason why a small subset of genes feature distinctive PROMPTs is unclear: there was no discernible difference in terms of sequence motifs previously associated with exosome sensitivity (Ntini et al., 2013; Almada et al., 2013), and no apparent shared role between genes having PROMPTs. Principally, the lack of detected PROMPTs may be either due to (i) absence of transcription initiation on the reverse strand upstream of *Arabidopsis* mRNA TSSs or (ii) extremely efficient degradation of PROMPTs by redundant RNA degradation systems, so that transcripts will not be detectable even when the exosome is rendered non-functional. The lack of signal in nascent RNA assays together with our failure to detect widespread occurrence of PROMPTs using double mutants constitute a strong argument for hypothesis (i), although it is difficult to entirely rule out hypothesis (ii), because not all relevant RNA degradation systems can be mutated simultaneously and because even GRO-seq signal may be lost through the action of nuclear degradation enzymes targeting nascent RNA.

Similarly, we found evidence of bidirectional transcription at intergenic NDRs producing exosome sensitive transcripts, which feature many of the same signatures as mammalian enhancer regions, including chromatin states. To our knowledge, this has previously not been reported in *Arabidopsis,* but it remains to be discovered whether these enhancer-like regions bind transcription factors and have a role in regulating distal gene transcription initiation.

### Possible reasons for PROMPT rarity in plants

Assuming that PROMPT initiation is rare, it is interesting to speculate how *Arabidopsis* promoters, unlike those of vertebrates, can initiate in only one direction and what might have favored the evolution of promoter directionality. A genetic study in yeast has shown that promoter directionality can be enforced by chromatin-based mechanisms: the histone chaperone, Chromatin Assembly Factor I (CAF-I) that facilitates replication-coupled nucleosome assembly (Smith and Stillman, 1989) restricts divergent transcription, while histone exchange promoted by acetylation of Lys56 on histone 3 (H3K56) promotes divergent transcription (Marquardt et al., 2014). It is possible that similar or other independently evolved chromatin-based mechanisms act in plants to enforce promoter directionality. One obvious reason for repression of PROMPT initiation in *Arabidopsis* may be the high gene-density of its genome: since genes are closely located, PROMPTs may interfere with upstream gene transcription. However, the fact that PROMPTs are also rare in maize with a less gene-dense and larger genome than many vertebrates (Erhard et al., 2015; Lozano et al., 2018) does not support this hypothesis and suggests that PROMPT suppression may be more general in plants. What might drive suppression of upstream transcripts specifically in plants within the eukaryotic kingdom? One provocative, if very speculative, possibility could be the fact that plants use the small RNA-dependent pathway RNA directed DNA methylation (RdDM) to drive DNA methylation and transcriptional repression to transposable elements and other repetitive regions of the genome. Maintenance of RdDM requires continued transcription of targeted loci by specialized RNA polymerases, but the initiating event in RdDM would logically require the presence of an RNA for targeting by small guide RNAs. Thus, we speculate that the prevalence of PROMPTs may be selected against in plants, as it would constitute a rich source of potential off-targets for small RNAs involved in RdDM and thereby jeopardize stringent transcriptional control of endogenous genes.

### A large class of exosome-sensitive antisense RNAs initiating in 3’ regions of protein-coding genes

The large number of cases of antisense TSSs residing in 3’-UTRs of mRNAs is an unexpected finding of our study, and distinct from earlier studies showing peaks of RNAPII or GRO-seq signal at or downstream of 3’ ends on the sense strand (e.g. (Zhu et al., 2018)). These TSSs define initiations of long, exosome-sensitive non-spliced antisense RNAs, which often cover large parts of the cognate mRNA. The function of these transcripts, if any, is unclear, although it is interesting to note a clear over-representation in genes encoding transcription factors: antisense transcripts may be repressing the mRNA by RNA-RNA hybridization, or the act of transcription of these RNAs may repress transcription of cognate mRNAs by steric effects such as RNAPII collision, or clearing of DNA-binding proteins and chromatin states. Indeed, there is already evidence for regulation of developmental genes by antisense RNAs, including the DOG1/asDOG1 example in Fig 8, which are very similar to the class of RNAs we show here to be prevalent.

## METHODS

### Plant materials

All *Arabidopsis thaliana* plants are of the Columbia ecotype (Col-0). The *hen2-4* (AT2G06990, SALK_091606C) and *lsm8-2* (AT1G65700, SALK_048010) mutants were described in (Lange et al., 2011) and (Perea-Resa et al., 2012). Seeds of these mutants were kind gifts from Dominique Gagliardi (*hen2-4*) and Julio Salinas (*lsm8-2*). The *rrp4-2* hypomorphic mutant is described in (Hématy et al., 2016) and was kindly provided by the authors.

### Genotyping

DNA was isolated by adding one volume of Phenol-Chloroform 50:50 to freshly ground leaves in urea buffer (42% (w/v) urea, 312.5 mM NaCl, 50mM Tris-HCl pH 8, 20 mM EDTA, 1% N-lauroylsarcosine). Phases were separated by centrifugation and the supernatant containing DNA was isolated. Nucleic acids were precipitated with one volume of isopropanol. DNA was pelleted by centrifugation and rinsed with EtOH 70%. The resulting purified DNA was used as a template for polymerase chain reaction to confirm the T-DNA insertion in *hen2-4* and *lsm8-2*. The G55E single-point mutation in the *rrp4-2* mutant was verified by target DNA amplification followed by enzymatic digestion (Eco47I, *AvaII*). Genotyping primers are available in **Supplementary Table S1**.

### Growth conditions and total RNA extraction

Wildtype, *hen2-4*, and *rrp4-2* seeds were surface-sterilized first by incubation in 70% EtOH (2 min) followed by 1.5% sodium hypochlorite and 0.05% Tween-20 (10 min), and rinsed three times with autoclaved ddH_2_O. Seeds were stratified in complete darkness at 4°C for 72 hours then germinated on 1% Murashige & Skoog (MS) media (4.4 g/L MS salts mixture, 10 g/L D-sucrose, 8 g/L agar) at pH 5.7 under sterile conditions and cycles of 16 hours light/8 hours of darkness. Intact 12 day-old seedlings were transferred in 8 mL of 1% liquid MS media (4.4 g/L MS salts mixture, 10 g/L D-sucrose) in 6 well-plates (Nunc) and allowed to acclimate for 48 hours with mild agitation (130 rpm).

Total RNA was extracted from a pool of 10 complete and intact 14 day-old seedlings. Plant materials were flash-frozen and 1 mL of TRI-Reagent (Sigma-Aldrich) was added to 100 mg of finely ground tissue and vortexed directly. Phase separation was achieved by adding 200 µL of chloroform, vigorous shaking, and centrifugation at 4°C (10 min, 15000 rpm). The aqueous phase was transferred to a fresh tube and one volume (400 µL) of isopropanol was added before RNA precipitation at room temperature (30 min). The total RNA was pelleted after centrifugation at 4°C (15 min, 15000 rpm) and rinsed three times with 70% EtOH prior to solubilization in autoclaved ddH_2_O. To rid the total RNA of contaminants and obtain higher quality material, polysaccharide precipitation was carried out using the double sodium acetate method. Total RNA was assessed for concentration and purity using NanoDrop ND-1000 (Thermo Scientific) and checked for degradation on Bioanalyzer 2100 (Agilent).

### CAGE libraries, filtering and mapping

CAGE libraries were prepared as in (Takahashi et al., 2012) with a starting material of 5 µg of total RNA. The National High-Throughput DNA Sequencing Centre of the University of Copenhagen performed the sequencing on an Illumina HiSeq 2000 platform. As recommended by Illumina, 30% of Phi-X spike-ins were added to each sequencing lane to balance the low complexity of the 5’ ends of the CAGE libraries. FASTX Toolkit v0.0.13 (http://hannonlab.cshl.edu/fastx_toolkit) was used to remove the linker sequences, retain the first 25 nt from the 5’ end, and to filter for a minimum quality of Q30 (Phred score) in 50% of the remaining bases. The clean reads were mapped on TAIR10 (Ensembl release 26) reference genome with Bowtie v1.1.2 (Langmead et al., 2009) using the parameters -t best -strata -v -k 10 -y -p 6 -phred33-quals -chunksmbs 512 -e 120 -q -un. Uniquely mapped CAGE tags 5’ ends were counted at each genomic position to obtain a set of CAGE transcription start sites, typically referred to as ‘CTSSs’ in the CAGE literature. CTSS coordinates were offset by 1 bp to account for G-addition bias (Carninci et al., 2006).

### Quantification and clustering of CAGE TCs

Most analyses were performed with the CAGEfightR v1.2 package (Thodberg et al., 2019) in the Bioconductor environment. Only CTSSs with at least one count in a minimum of 3 libraries (the smallest sample group) were kept. CTSSs were normalized into tags-per-million (TPM) to mitigate differences in sequencing depths. A pan-experiment set of CAGE Tag Clusters (TCs) was established by computing the sum of TPM-values over all libraries at each base pair, followed by neighbour-clustering of CTSSs on the same strand (within a distance of 20 bp) (**Supplementary Data S4**). Only TCs supported in at least 2 libraries with a pan-experiment minimum of 1 TPM were considered for further analysis. TC peaks were defined as the position with maximum signal within the TC.

### Annotation of CAGE TCs

TAIR10 annotation was recovered from the TxDb.Athaliana.BioMart.plantsmart28 Bioconductor package. ARAPORT11 GFF3 annotation (June 2016) was obtained from www.araport.org and converted into a TxDb object (Lawrence et al., 2013) using the makeTxDbFromGFF function. Using TC peaks as proxy (see above), annotation of CAGE TCs was conducted against a hierarchical annotation built from TAIR10 and ARAPORT11 individually (**see** Figure 1). The promoter region was defined as the 200 bp window centered on the annotated gene TSS. The promoter-proximal region extends 400 bp upstream of the annotated TSS. The same regions, but on the opposite strand, was referred to as the reverse region. The antisense category covered the whole gene body but on the opposite strand. The 5’-UTRs, 3’-UTRs, and CDS regions were defined as in TAIR10 or ARAPORT11. The exonic category indicate non-protein-coding exons, including exons of long non-coding RNAs. In case of multiple overlaps, a CAGE TC peak was annotated using the highest-priority category of the overlapping annotations in the annotation hierarchy.

### Alternative TSS analysis

CAGE TCs attributed to a TAIR10 annotated gene, and contributing at least 10% of the total TPM gene expression across the gene, were selected as a basis for assessing the landscape of alternative TSSs in unchallenged wild type seedlings.

### Recall of TSSs/TCs across datasets

TCs and TSSs from CAGE wild type, PEAT-seq, and nanoPARE datasets were merged. To obtain a non-redundant set of transcription start sites, overlapping regions were collapsed. The number of recalled TCs/TSSs from the non-redundant set was computed for each dataset and plotted as a Venn diagram (Figure 3A).

### Average meta-profiles

All metaplots were computed on normalized read counts using the TeMPO package (https://github.com/MalteThodberg/TeMPO) with trimmed signal (0.01% to 0.99% percentiles).

### Quantification and comparison of TSS across datasets

TSSs from ARAPORT11, TAIR10, nanoPARE and PEAT-seq were extended by 100 bp in both directions and quantified using the CAGE wt TPM signal. To account for the large differences in expression as observed in Figure 3B, TSSs were filtered to have at least 1 CAGE TPM in at least 2 samples and were subsequently used as anchors for comparison of nascent RNA signals, MNase signal, and sequence patterns.

### RNA-seq libraries, mapping and quantification

Seeds for wt, *lsm8-2*, *rrp4-2,* and the *rrp4-2 lsm8-2* double mutant were subjected to the same sterilization, stratification, and growth conditions as described for the CAGE samples. Total RNA was purified from complete and intact 14 day-old seedlings and polysaccharide precipitation was used as for the CAGE samples, see above. The resulting polysaccharide-free total RNA was sequenced by Novogene (Hong-Kong) as stranded Illumina paired-end 150 bp reads. Libraries underwent ribosomal depletion (Ribo-zero) and the fragment size selection limit was lowered to 200 nt to allow for the capture of hypothetical PROMPT transcripts. Adapters were removed from the raw reads using Cutadapt v2.3 (Martin, 2011). Mapping to the TAIR10 reference genome (Ensembl release 26) was processed with STAR v2.7.0 (Dobin and Gingeras, 2015), only considering unique mappers and concordant pairs. The matrix of counts antisense to TAIR10 annotated genes was generated with featureCounts v1.6.3 (Liao et al., 2014) with parameters -O --minoverlap 3 --largestOverlap -B -p -s 1 (**Supplementary Data S5**).

### DHS annotation

DHS regions, DNase-Seq and MNase-Seq were obtained from PlantDHS.org (Zhang et al., 2016). Position of the highest DNase signal in flower tissue was used to identify the DHS summit (DHSS) in each DHS region. DHSSs were extended 400 bp on both directions and annotated with TAIR10 reference, using the hierarchical strategy as defined in Figure 1C.

### Poly(A), 5’ and 3’ splice sites at mRNA & PROMPT TSSs

The seqPattern Bioconductor package was used to scan for AWTAA as well as 5’ and 3’ splice sites. For AWTAA, the consensus sequence was used. Position frequency matrices from (Brown et al., 1996) were used with a matching score threshold of 80% for the 5’ and 3’ splice sites.

### Differential expression analysis and GO-term enrichment

The edgeR (Robinson et al., 2010) and limma (Ritchie et al., 2015) packages were used to conduct differential expression analyses. Counts were normalized using the weighted trimmed mean of M-values (TMM) method (Robinson and Oshlack, 2010) and transformed using voom (Law et al., 2014) to model the mean-variance relationship prior to linear modeling. Resulting P-values were corrected for multiple testing with the Benjamini-Hochberg method. For gene set enrichment analyses, the gProfileR package was used with the complete set of detected genes as the background universe.

### Support of PROMPTs by nascent RNA sequencing

Average signal of 5’ GRO-cap (Hetzel et al., 2016), as well as GRO-seq from two labs (Hetzel et al., 2016; Liu et al., 2018), was calculated across the genome-wide PROMPT regions, defined as the −400 bp antisense stretches from TAIR10 annotated TSSs. Support of exosome-sensitive PROMPT regions by nascent sequencings was considered upon showing 2-fold the average genome-wide signal.

### *De novo* reconstruction of antisense transcripts

To assemble *de novo* transcripts from RNA-seq data, Cufflinks v2.2.1 (Roberts et al., 2011) was used with default parameters, with the TAIR10 general feature format (GFF) annotation provided as a guide (option -g). Cuffcompare was applied to compare the set of *de novo* transcripts across genotypes. *De novo* transcripts overlapping a reference exon or intron but on the opposite strand (codes s and x, respectively), and associated with an RNA-seq up-regulated gene on the reverse strand were selected for assessing the length and splicing of exosome-sensitive transcripts.

## Author contributions

AT conducted experiments. JL constructed the CAGE libraries. AT and MI did the computational analyses. AT made all figures. AT, PB and AS interpreted the results. AT, AS, PB wrote the paper with inputs from all authors.

## Acknowledgements

Kian Hématy (*rrp4-2*), Julio Salinas (*lsm8-2*) and Dominique Gagliardi (*hen2-4*) are thanked for providing mutant seeds. We thank Michael Schon for critical assessment of nanoPARE genomic alignments. Work in AS lab was supported by grants from Novo Nordisk Foundation and Lundbeck Foundation. Work in PB lab was supported by grants from Novo Nordisk Foundation (Hallas Møller stipend 2010), and Villum Foundation (project grant 13397).

---

**Supplementary Figure 1.**
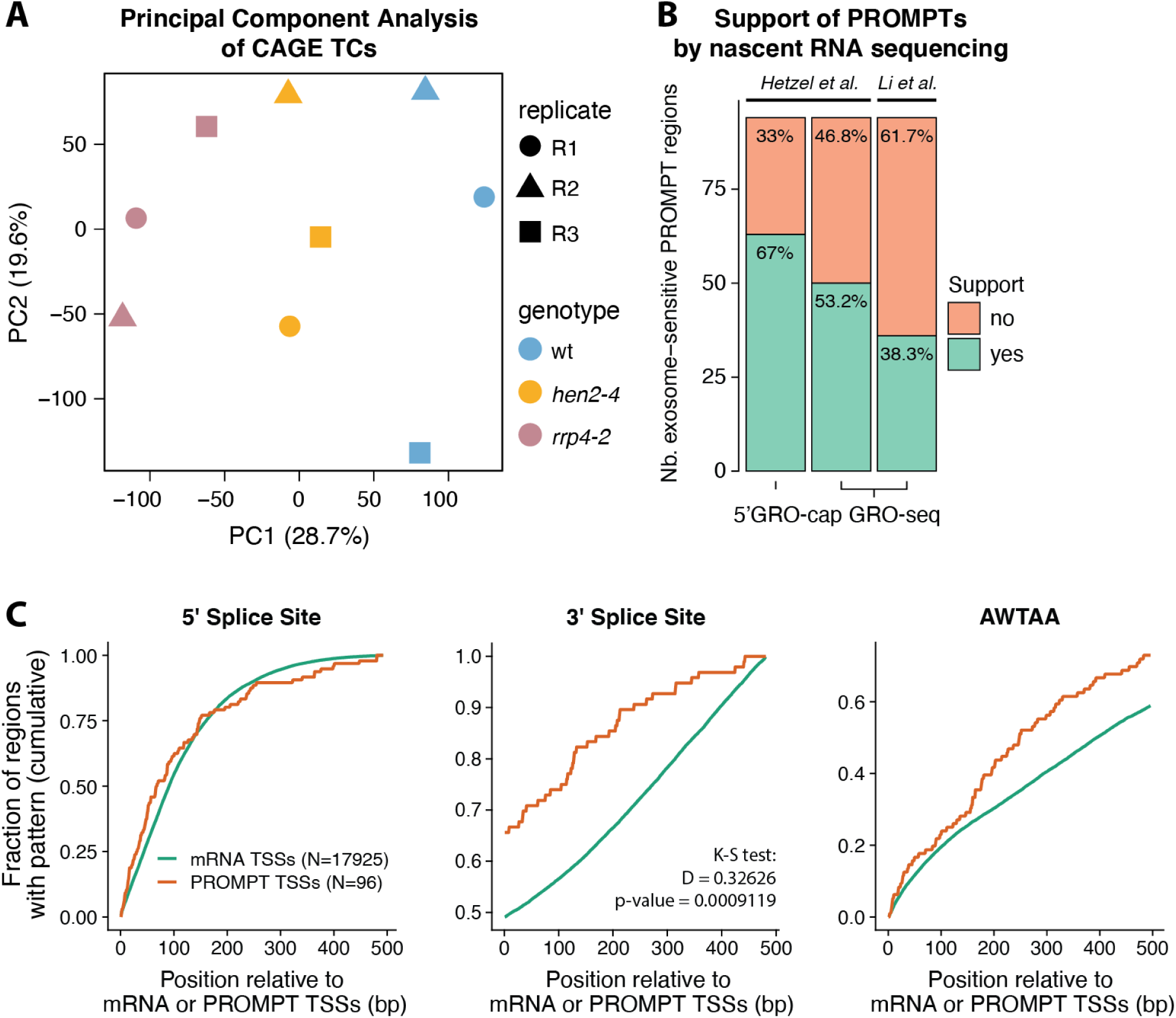
Principal component analysis and characterization of PROMPT regions. **(A) Principal component analysis of CAGE TCs.** Axes show the two first principal components and the percentage of explained variance. Each point represents a CAGE library, whose genotype and replicate are indicated by colors and shapes, respectively. **(B) Support of PROMPTs by nascent RNA sequencing data.** X-axis shows 5’ GRO-cap and GRO-seq from two laboratories, as indicated on top. Y-axis counts the number of exosome-sensitive PROMPT regions detected in this study, colored by whether they were supported by nascent transcription data or not (see Methods). **(C) AWTAA poly(A) and splice sites at mRNA and PROMPT TSSs.** X-axis shows distance relative to CAGE TC peaks associated with either mRNA promoters or exosome-sensitive PROMPTs, in bp. Y-axis shows the fraction of regions with a 5’ or 3’ splice site (left and middle) or AWTAA poly(A) (right) pattern.

**Supplementary Figure 2.**
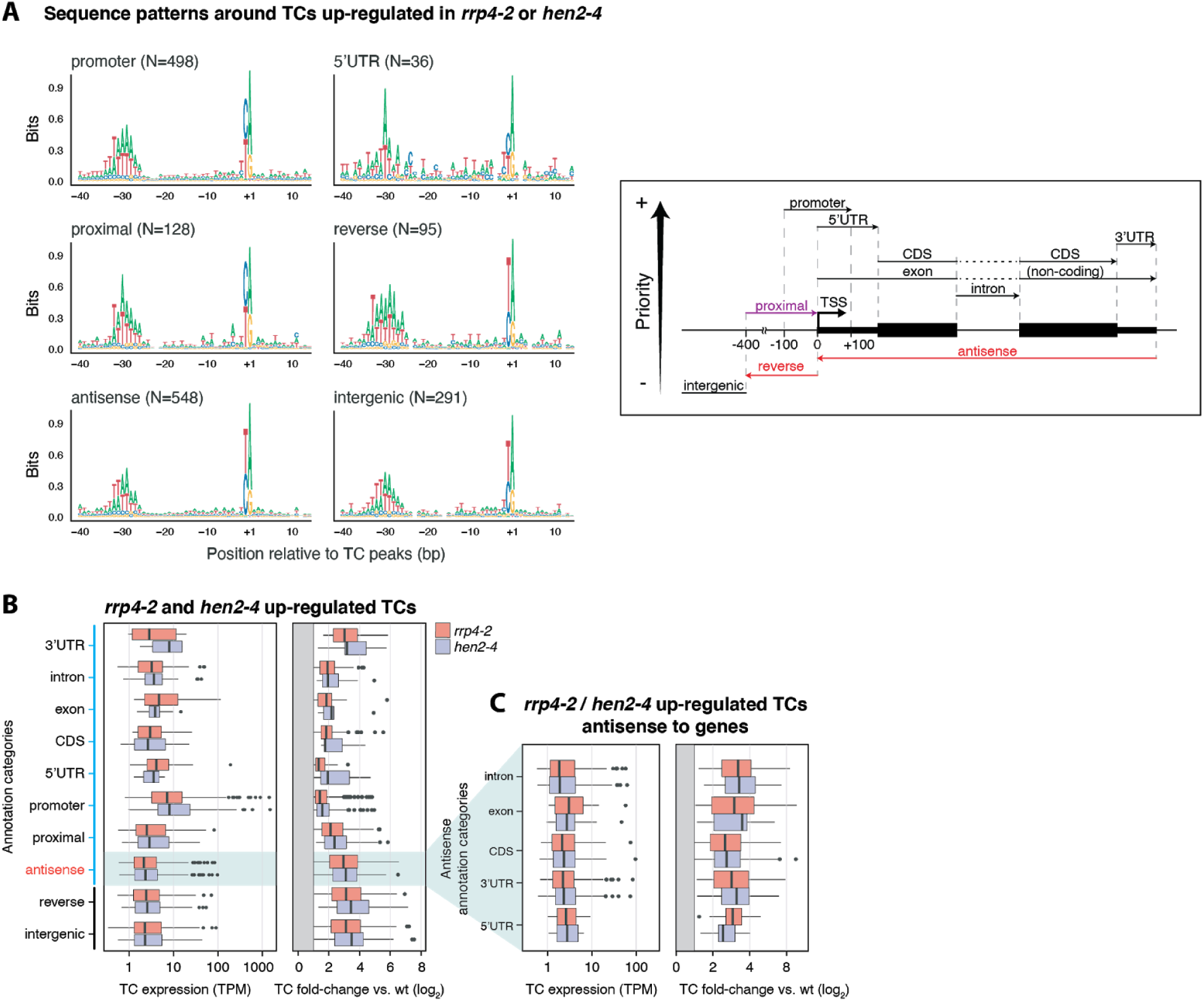
Sequence and expression properties of exosome-sensitive TCs. **(A)Sequence patterns of CAGE TCs up-regulated in *hen2-4* or *rrp4-2*.** Organized as in Figure 3C. The annotation hierarchy strategy is recapitulated on the right side. **(B) Annotation, expression, and fold-change of up-regulated CAGE TCs.** Left panel: X-axis shows the expression of CAGE TCs in TPM. Y-axis distinguishes annotation categories based on TAIR10. Colors separate CAGE TCs up-regulated either in *rrp4-2 vs.* wt (red) or *hen2-4 vs.* wt (violet). Right panel: X-axis shows log_2_ fold-change of respective mutant *vs.* wt. **(C) Expression and fold-change of up-regulated CAGE TCs, antisense to a gene.** Left and right panel organized as in B, but shows expression of CAGE TCs that are antisense to an annotated TAIR10 gene.

**Supplementary Figure 3.**
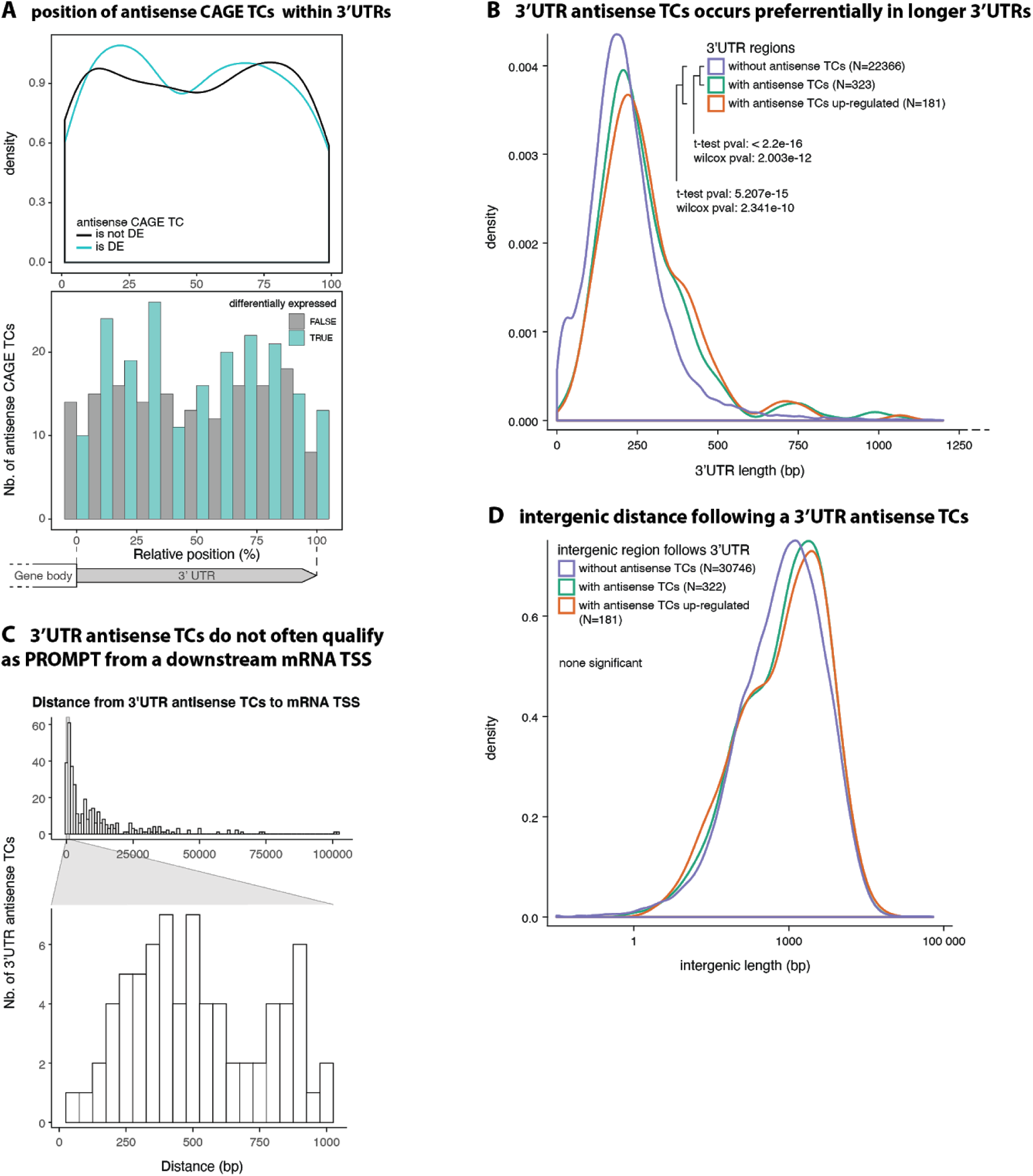
Investigation of CAGE TCs antisense to an annotated 3’-UTR. **(A) Positioning of antisense TCs within 3’-UTRs.** X-axis show position within 3’-UTR, scaled from 0 (start of UTR) to 3’-end (100). Top panel shows the density of TCs along x, colored by whether TCs are up regulated (is DE) in *rrp4-2 vs.* wt or not (not DE). Lower panel shows the same data as bar plots, counting the number of cases within X-axis bins. **(B) Relation between 3’-UTR length and presence of antisense TCs.** Density distribution of 3’-UTR lengths, where colors indicate the presence of antisense TCs and their up-regulation status. **(C) 3-’UTR-antisense TCs are not PROMPTs of neighboring promoters.** X-axis shows the distance between 3’-UTR-antisense TC peaks and the closest annotated mRNA TSS, located downstream of the PROMPT and on the opposite strand, in bp. Y-axis indicates the number of 3’-UTR-antisense TCs. Bottom insert zooms on distances below 1000 bp. **(D) Relation between intergenic gene distance and presence of antisense TCs.** Density distribution of lengths intergenic regions (to next annotated gene) downstream of an annotated 3’-UTR. Colors indicate the presence of antisense TC in the 3’-UTR and up-regulation status.

## List of Supplementary Tables and Data sets

### Supplementary Table

S1: Genotyping primers

### Supplementary Data Sets

S1: Unidirectional and bidirectional CAGE TCs, and PROMPTs

S2: Results of differential expression analyses

S3: RNA-seq *de novo* reconstructed transcripts

S4: expression matrix of CAGE TCs (TPM-normalized)

S5: expression matrix of RNA-seq (raw counts, antisense to annotated genes)

## Notes

#### Summary of Updates

We found an alignment 1 bp indexing error on minus strand for nanoPARE TSS data (Thanks to Michael Schon for making us aware of the issue), which is used for comparison with CAGE data in Figure 3. This is now corrected. This changes changes the nanoPARE logo in Figure 3C and very subtle the nanoPARE profile plots in Figure 3. Small text changes are made to comment on the change. This does not change any major conclusions in the paper.

