## Supplementary table 1 for "Characterization of *Arabidopsis thaliana* promoter bidirectionality and antisense RNAs by depletion of nuclear RNA decay enzymes"

| **Genotype** | **AGI** | **Left primer 5'-3'** | **Right primer 5'-3'** | **Comment** |
| --- | --- | --- | --- | --- |
| *hen2-4* | AT2G06990 | TATGGTATTCAGCAACCTCCG | GTTCCTCAAATGCTGCTCTTG | SALK_091606 genotyping |
| *lsm8-2* | AT1G65700 | ACTAACTGGCCTCTGAATGGAAG | AAGAAGACCCAAGACTCCGATG | SALK_048010 genotyping |
| *rrp4-2 (sop2-1)* | AT1G03360 | CTATTCCCGTCAACCATGACG | CATCGACCTCGGAAGTTCCAGGT | DNA amplification before enzymatic digestion |
| */* | / | ATTTTGCCGATTTCGGAAC | / | SALK Right Border primer LBb1.3 |
